# Humans reconfigure target and distractor processing to address distinct task demands

**DOI:** 10.1101/2021.09.08.459546

**Authors:** Harrison Ritz, Amitai Shenhav

## Abstract

When faced with distraction, we can focus more on goal-relevant information (targets) or focus less goal-conflicting information (distractors). How people use cognitive control to distribute attention across targets and distractors remains unclear. To help address this question, we developed a parametric attentional control task that can index both target discriminability and distractor interference. We find that participants exert independent control over target and distractor processing. We measured control adjustments through the influence of incentives and previous conflict on target and distractor sensitivity, finding that these have dissociable influences on control. Whereas incentives preferentially led to target enhancement, conflict on the previous trial preferentially led to distractor suppression. These distinct drivers of control altered sensitivity to targets and distractors early in the trial, promptly followed by reactive reconfiguration towards task-appropriate feature sensitivity. Finally, we provide a process-level account of these findings by showing that these control adjustments are well-captured by an evidence accumulation model with attractor dynamics over feature weights. These results help establish a process-level account of control reconfiguration that provides new insights into how multivariate attentional signals are optimized to achieve task goals.

## Introduction

Whether we are having a conversation in a crowded coffee shop or writing a paper at our desk while surrounded by browser tabs, most tasks require us to engage in two distinct forms of attentional control. One form of control enhances the processing of task-*relevant* information, for instance by paying careful attention to what our conversation partner is sharing with us. The other form of control suppresses the processing of task-*irrelevant* information, particularly that which conflicts with our primary goal (e.g., distraction from a nearby conversation). While past research has extensively studied target and distractor processing, it has done so primarily by focusing on each one separately. As a result, relatively little is known about how control over task-relevant information (targets) interacts with control over task-irrelevant information (distractors), and under what situations one would be prioritized over the other. Here, we bridge previous methodological gaps to gain new insight into how control is differentially allocated to achieve these aims, and provide an integrative model that accounts for the dynamics by which these two forms of control are adjusted within and across trials.

Research into how people enhance the target of their attention versus actively suppress distractors has been largely governed by separate research areas, using different approaches. Studies of perceptual decision-making have characterized the process by which people try to determine the correct response (e.g., which of two categories this stimulus belongs to) based on noisy information about a target stimulus, and how this varies with the difficulty of discriminating that stimulus (e.g., how perceptually similar two stimuli are; Britten et al., 1992; Gold and Shadlen, 2007). This contrasts with studies of inhibitory control, in which the correct response to a target is typically unambiguous (e.g., respond left when seeing a high-contrast leftward-facing arrow), but a second dimension of the stimulus display (one that is typically processed more automatically; e.g., flanking arrows pointing rightward) triggers a conflicting response (Botvinick and Cohen, 2014; Posner and Snyder, 1975).

Despite the substantial progress that has been made in understanding these two processes in parallel, critical questions remain that can only be addressed by studying them in tandem (Ritz et al., 2022). Most notably, it is unclear how people decide how to distribute their control between targets and distractors. When the demands or incentives for performing a task change, do people re-direct control towards target enhancement, distractor suppression, or both? For instance, previous work has shown that people are less susceptible to the influence of distractors after overcoming a previously conflicting distractor (the so-called *conflict adaptation* or *congruency sequence* effect; Gratton et al., 1992). Prevailing models have accounted for these findings by assuming that participants increase attention to the target dimension following a high-conflict trial (Botvinick et al., 2001; Egner, 2007), but limitations of the relevant experiments (in particular, that their focus on variability in distractor congruency) make it difficult to rule out that adaptation is occurring at the level of distractor suppression instead (Lindsay and Jacoby, 1994; Tzelgov et al., 1992). It is more generally an open question whether effects of recent task difficulty (e.g., low discriminability or high-conflict) result in control-specific or control-general adaptations and, similarly, whether the motivation to improve performance in such settings leads to preferential engagement of one or both forms of control.

One way that previous research has studied target and distractor adjustments is to measure changes in brain activity associated with task-relevant stimulus processing. For example, some previous work has suggested that conflict-triggered control preferentially enhances sensitivity in regions associated with target stimuli (e.g., faces in fusiform face gyrus; Egner and Hirsch, 2005). Other studies have found evidence for both target and distractor control by using similar stimulus-tagging methods (Gazzaley et al., 2005; Soutschek et al., 2015) or by exploiting lateralized EEG responses (Noonan et al., 2016; Wöstmann et al., 2019). The range of results across these neuroimaging experiments may come from the different tasks that have been deployed (Egner, 2008), but may also arise from noisy or complex correspondence between neuroimaging methods and underlying cognitive processes. In the current experiment, we provide new methods for indexing target and distractor sensitivity from behavior alone, enabling us to provide new insight into the cognitive architecture of feature-selective control.

Recent models of controlled decision-making have emphasized the role that within-trial attentional dynamics play in response conflict tasks, offering new insight into the implementation of cognitive control (Servant et al., 2014; Weichart et al., 2020; White et al., 2011; Yu et al., 2009). These models have largely focused on the Eriksen flanker task, modelling how an attentional spotlight centered on the target item narrows over time. This formulation necessarily yokes target enhancement and distractor suppression due to the spatial spread of attention. As a result, little is known about whether target and distractor processing dynamics can fall under independent control when these are not explicitly yoked, as in the case of feature-based attention. Less still is known about how adjustments driven by factors like conflict adaptation and incentives alter the *dynamics* of target and distractor processing (Adkins and Lee, 2021).

To address these questions, we developed a novel task that orthogonally varies target and distractor information, measuring how processing of these two dimensions varies both within and across trials. Our task merges elements of paradigms that have been separately popularized within the two research areas above. To capture variability in target processing, we based our task on the random dot kinematogram paradigm (Danielmeier et al., 2011; Kang et al., 2021; Mante et al., 2013; Shenhav et al., 2018). This task parametrically varies the motion discriminability (e.g., percentage of dots moving left) and color discriminability (e.g., percentage of green dots) across an array of dots. Participants were instructed to respond to the color dimension, while ignoring the motion dimension. Critically, whereas color response mappings were arbitrary (e.g., left hand for green), motion responses were exactly aligned with the direction of motion (e.g., left hand for leftward moving stimuli), resulting in potent “Simon-like” response interference from this prepotent distractor.

Previous work has demonstrated response conflict and trial-to-trial adjustments in a color-motion kinematogram with full target coherence and binary distractor congruence (Danielmeier et al., 2011). We extended this task by parametrizing both target coherence and distractor congruence. In doing so, we are able to not only obtain more precise measures of feature sensitivity by accounting for global performance factors (e.g., lapse rate; Wichmann and Hill, 2001) but, importantly, to isolate how participants simultaneously configure attention towards each of these features. We are able to then further examine how these multivariate control configurations (i.e., joint setting of target and distractor gain) dynamically adjust to accommodate incentives and demands associated with task performance. We use the precision afforded by these methodological advances to inform an explicit process model of multivariate attentional control.

We find that participants independently and dynamically control target and distractor processing over the course of a trial. Participants preferentially enhance target sensitivity under incentives, and preferentially suppress distractor sensitivity after high conflict trials. Moreover, they implement these control strategies by changing the initial conditions of a dynamic process that enhances task-relevant feature processing and suppresses task-irrelevant feature processing.

Finally, we find that these control strategies can be captured by extending classic neural network models of cognitive control to incorporate an attractor network that dynamically regulates the influence of different task features on choice. Together, these results extend our understanding of both decision-making and cognitive control by bridging the methodological and theoretical divides between these fields, providing new assays and process models for the top-down control of information processing.

## Results

Participants performed the Parametric Attentional Control Task (PACT), a perceptual discrimination task that required them to classify the dominant color in an array of moving dots (Figure 1a). Participants made bimanual responses, for example responding with their left hand when the dominant color was purple or blue or responding with their right hand when the dominant color was green or beige. To avoid stimulus repetition priming (Braem et al., 2019), two colors were assigned to each response and the majority color did not repeat across sequential trials. Across trials, we varied the extent to which those dots were coherently moving in the same or opposite direction as the correct response (distractor interference; Experiments 1-3) and how easily the participant could determine the dominant color (target discriminability; Experiments 2- 3; Figure 1b). Participants performed the main *Attend-Color* PACT in blocks of 100 trials. To enhance the potency of motion as a distracting dimension (Shiffrin and Schneider, 1977) and allow for additional measures of automaticity and feature-specificity, participants alternated between these blocks-of-interest and shorter blocks (20-50 trials) in which participants instead responded to the direction of dot motion (*Attend-Motion* PACT; Figure 1c).

**Figure 1.**
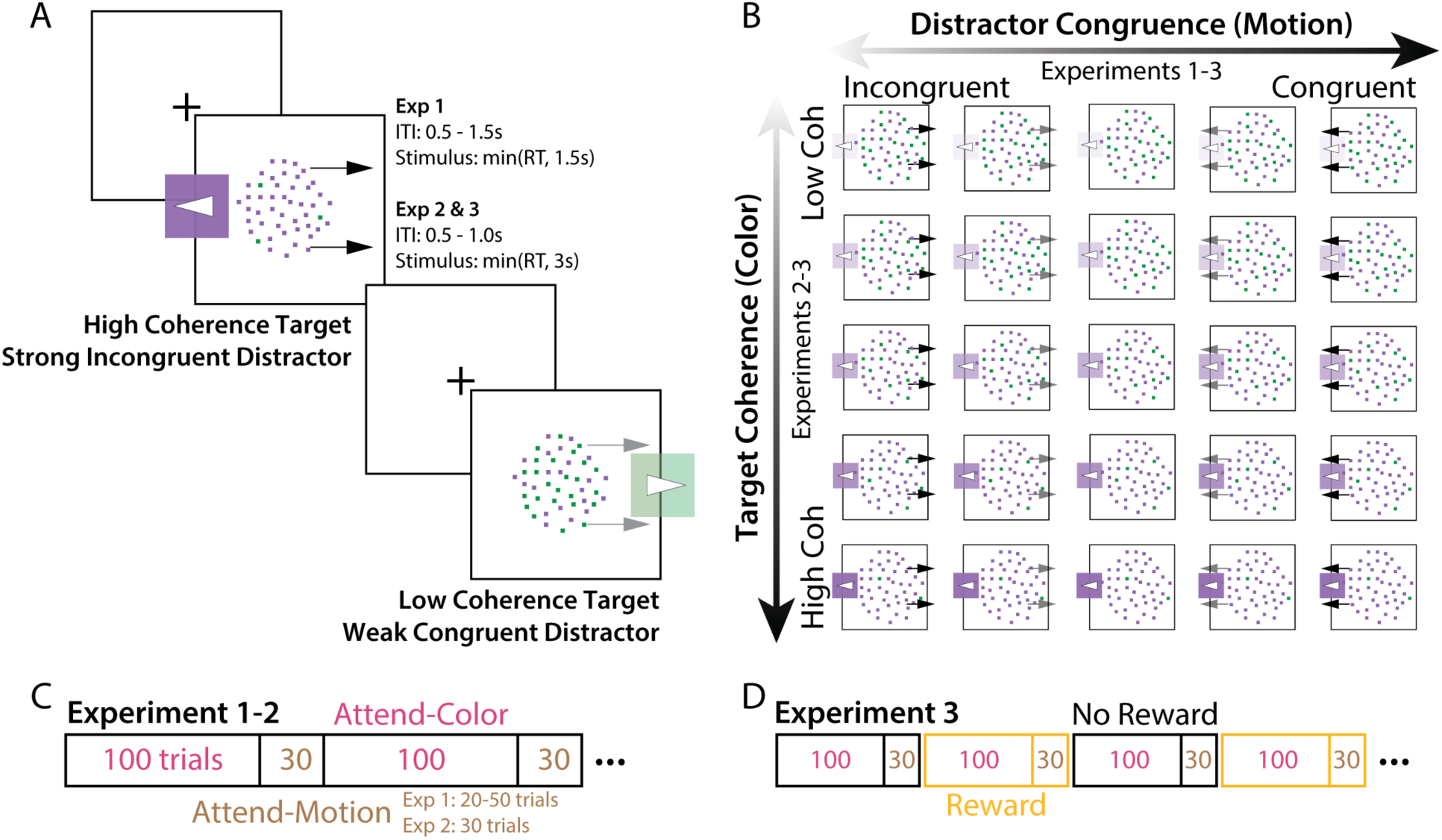
*Parametric Attentional Control Task (PACT).* **A)** On each trial, participants responded to the dominant color in a bivalent random dot kinematogram. This stimulus had a random color (target) coherence, depending on the proportion of dots that were in the majority. This stimulus also had a random motion (distractor) congruence, depending on motion coherence in the same or opposite direction as the color response. **B)** Across trials, we parametrically and independently varied the coherence of the dominant color (y-axis) and the congruence of the motion direction (x-axis). **C)** In Experiments 1 and 2, participants alternated between longer blocks of Attend-Color trials (target dimension was color, as in A) and shorter blocks of Attend-Motion trials (target dimension was motion). Participants took a self-timed break between blocks. **D)** In Experiment 3, participants alternated between pairs of Reward blocks and No Reward blocks. On Reward blocks, participants could earn a monetary bonus if they were fast and accurate, whereas we just encourage good performance on No Reward blocks. Participants were informed of the reward condition during their break between blocks.

### Task performance varies parametrically with target discriminability and distractor interference

In Experiment 1 (N = 56), participants performed the PACT with uniformly colored dots (e.g., all blue or all green), but with the dots moving in a direction either congruent or incongruent with that target response. We varied the strength of this distractor dimension between being fully congruent with the correct color response (100% leftward coherence for a left color response) to being fully incongruent (100% rightward coherence for a left color response; Figure 1b). For trials mid-way between these two extremes (cf. ‘neutral’ trials), the dots did not move consistently in one direction or another (0% motion coherence).

Consistent with past research on cognitive control, we found that participants were slowest and least accurate when distractors were fully incongruent (median RT = 585ms, mean accuracy = 89%) and fastest and most accurate were fully congruent (median RT = 553ms, mean accuracy = 97%; cf. (Danielmeier et al., 2011). Performance on neutral trials (0% motion coherence) fell between these two extremes (median RT = 576ms, mean accuracy = 94%). Extending this work, hierarchical regression analyses (see Methods) revealed that performance varied in a graded fashion across this continuum of interference. Both accuracy (Cohen’s *d* on regression estimate; *d* = -1.47) and reaction time (*d* = 1.25) worsened with parametrically increasing levels of interference (*p*s < 0.001, Figure 2c, Table 1).

**Figure 2.**
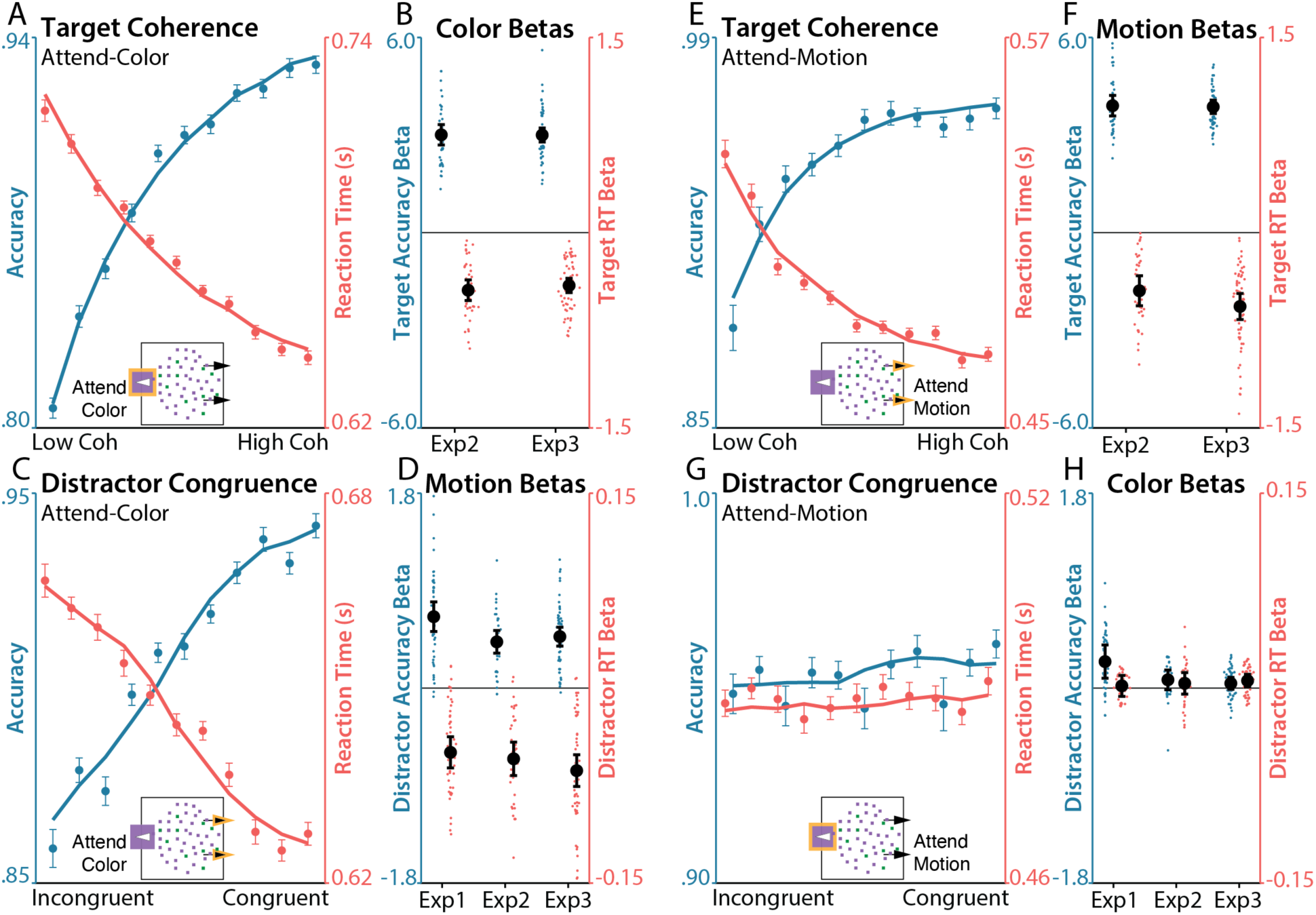
*Target and distractor sensitivity.* **A)** Participants were more accurate (blue, left axis) and responded faster (red, right axis) when the target color had higher coherence. Lines depict aggregated regression predictions. In all graphs, behavior and regression predictions are averaged over participants and experiments. Data aggregated across Experiments 2 & 3. **B)** Regression estimates for the effect of target coherence on performance within each experiment, plotted for accuracy (blue, left axis) and RT (red, right axis). **C)** Participants were more accurate and responded faster when the distracting motion had higher congruence (coherence signed relative to target response). In all graphs, behavior and regression predictions are averaged over participants and experiments. Data aggregated across Experiments 1-3. **D)** Regression estimates for the effect of distractor congruence on performance within each experiment, plotted for accuracy and RT. **E-F)** Similar to A-B, performance (E) and regression estimates (F) for the effects of target coherence during Attend-Motion blocks, in which motion was the target dimension. **G-H)** Similar to A-B, performance (G) and regression estimates (H) for the effects of distractor congruence during Attend-Motion blocks, in which color was the distractor dimension. Error bars on behavior reflect within-participant SEM, error bars on regression coefficients reflect 95% CI. Psychometric functions are jittered on the x-axis for ease of visualization.

**Table 1.** Target and distractor sensitivity.

| Table 1. Target and distractor sensitivity |  |  |  |  |  |
| --- | --- | --- | --- | --- | --- |
| DV | Predictors | Exp 1 (df = 45)<br>Effect size ($d$) | Exp 2 (df = 25)<br>Effect size ($d$) | Exp 3 (df = 45)<br>Effect size ($d$) | Aggregate<br>$p$-value |
| Choice | Target coherence |  | 3.27 | 3.73 | **$1.01 \times 10^{-44}$** |
|  | Distractor congruence | 1.47 | 1.42 | 1.50 | **$4.89 \times 10^{-32}$** |
| | Target × Distractor | | -0.184 | -0.344 | **0.0226** |
| RT | Target coherence |  | -1.90 | -1.99 | **$1.59 \times 10^{-28}$** |
|  | Distractor congruence | -1.25 | -1.49 | -1.43 | **$1.26 \times 10^{-29}$** |
| | Target × Distractor | | 0.230 | 0.0525 | 0.437 |

Effect sizes are calculated from MAP group-level regression estimates. $P$ -values are aggregated across experiments, with statistically significant $p$ -values (two-tailed, $\alpha = 0.05$ ) shown in bold.

In Experiment 2 (N = 40) and Experiment 3 (N = 60), participants performed the same task as in Experiment 1, but we additionally varied the discriminability of the target (color) dimension.

Across trials, the proportion of the majority color (*color coherence*) varied parametrically to make color discrimination easier (higher coherence) or more difficult (lower coherence). As in Experiment 1, the level of motion interference also varied across trials, independently of targets.

Consistent with past research on perceptual decision-making (Britten et al., 1992; Mante et al., 2013), we found that discrimination performance improved with higher levels of target discriminability. Participants in both studies were faster (Exp 2: *d* = -1.90, Exp 3: *d* = -1.99) and more accurate (Exp 2: *d* = 3.27, Exp 3: *d* = 3.73) with parametrically increasing levels of color coherence (aggregate *p*s < 0.001; Figure 2a, Table 1). At the same time, we continued to find that participants were slower and less accurate when the goal-irrelevant movement of those dots was increasingly incongruent with the correct color response (see Figure 2b, Table 1).

Performance on our task varied parametrically with both color coherence and motion coherence, but these two coherence manipulations were designed to exert their influence on performance in different ways. Whereas variability in color coherence was intended to influence the stimulus uncertainty directly relevant to goal-directed decision-making (i.e., determining which response is the correct one), motion coherence was intended to exert a more automatic influence on response selection by facilitating responses consistent with the direction of motion. We confirmed this assumption regarding the relative automaticity of motion versus color processing by having participants perform interleaved blocks in which they responded based on motion and ignored color (‘Attend-Motion’). We found that participants were more sensitive to the now-relevant motion coherence, but were no longer sensitive to the now-irrelevant color congruence (Supplementary Table 1-2). This asymmetry suggests that participants’ decisions were not solely driven by the bottom-up salience of these features, as participants were more sensitive to color when it was relevant and less sensitive to motion when it was irrelevant, reflecting differential engagement of top-down control across the two tasks (Cohen et al., 1992).

### Target discrimination and distractor interference occur in parallel

We found that participants’ task performance varied parametrically with both the target discriminability and distractor congruence, both for choice and reaction time. We next sought to further understand the relationships between these changes in performance, within and across features.

First, we tested whether a given feature exerted a similar influence on both accuracy and RT. We found that this was indeed the case, as there was a significant correlation between the effect distractors had on accuracy and RT (*r*s < -0.87, *p*s < 0.001). The influences of target discriminability on accuracy and RT were also significantly correlated (*r*s < -0.54, *p*s < 0.001; Supplementary Table 3). Thus, participants who became faster with higher levels of a given feature’s strength also became more accurate, suggesting that accuracy and RT shared a common underlying process (e.g., evidence accumulation rate, which we return to below).

Second, we tested whether the influences of target discriminability and distractor congruence on performance were independent (e.g., distractors and targets are processed in parallel; (Lindsay and Jacoby, 1994; Servant et al., 2014) or instead modulatory (e.g., distractor congruence influences target sensitivity). If the two forms of feature processing modulated one another, we would predict that target and distractor coherence would interact in predicting performance. We did not find such an interaction in RTs (*d*s = 0.05 to 0.23, *p* = 0.33; Table 1), though we did find a small but significant interaction between target and distractor coherence on accuracy (*d*s = - 0.18 to -0.34, *p* = 0.023). For both studies, removing target-distractor interactions as predictors in our accuracy regressions improved model fit (Protected exceedance probability on AIC: Exp 2 PXP = 1; Exp 3 PXP = 1). If distractors had an antagonistic influence on target processing, we would also predict that target and distractor sensitivity would be negatively correlated across subjects. Contrary to this prediction, these effects were either not significantly correlated or positively correlated, both for RT (Exp 2: *r*(25) = .14, *p* = .48; Exp 3: *r*(45) = .44, *p* = .0019) and accuracy (Exp 2: *r*(25) = -.15, *p* = .45; Exp 3: *r*(45) = .12, *p* = .43), suggesting that individual differences in target and distractor processing were not antagonistic.

### Previous conflict preferentially suppresses distractor sensitivity

Within a given trial, we found that performance varies parametrically and independently with the coherence of target and distractor features. We next sought to understand how participants adapted their information processing *across* trials, to provide insight into the control processes that guide performance in this task. We measured how participants’ feature sensitivity changed after difficult (e.g., more incongruent) trials, an index of cognitive control known as conflict adaptation (Egner, 2007; Gratton et al., 1992). The classic effect is that participants show weaker congruence effects after incongruent trials than after congruent trials, with the traditional interpretation being that this reflects upregulated target sensitivity (Botvinick et al., 2001; Egner, 2007). Our task allowed us to build on this work to test whether this adaptation effect varies parametrically with distractor congruence. Critically, we can also test whether adaptation occurs through an influence of previous conflict on subsequent target enhancement, distractor suppression, or both. Finally, we can further test whether adaptation occurs due to the discriminability of the *target* on the previous trial.

Across all three of our studies, we found that participants’ sensitivity to the distractor dimension was robustly and parametrically influenced by the distractor congruence on the previous trial, as reflected both in their choice (*d*s = 1.44 to 1.74, *p* < .001; Figure 3a) and RT (*d*s = 0.83 to 1.79, *p* < .001; Figure 3b; Table 2). When the previous trial had congruent distractors, participants had strong sensitivity to the distractor congruence (Figure 3a-b, navy). When the previous trial had incongruent distractors, participants were much less sensitive to distractors (Figure 3a-b, red). These patterns are consistent with those typically observed in studies of conflict adaptation (Danielmeier et al., 2011; Egner, 2007), and further demonstrate gradations within these classic effects.

**Figure 3.**
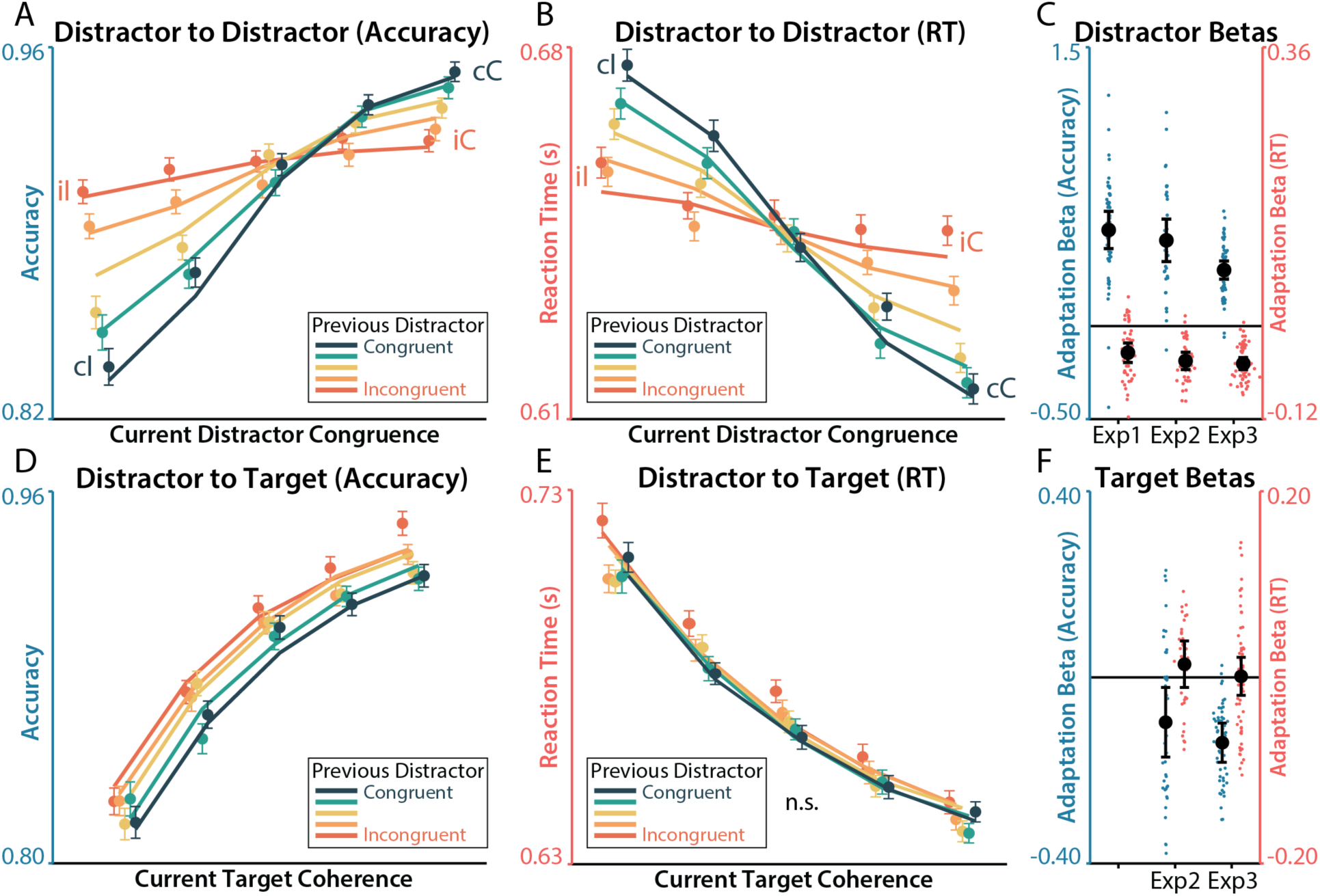
*Distractor-dependent adaptation.* **A-B)** The relationship between distractor congruence and accuracy (A) and RT (B) was weaker when the previous trial was more incongruent (redder colors). Lines depict aggregated regression predictions. **C)** Regression estimates for the current distractor congruence by previous distractor congruence interaction, within each experiment. **D-E)** The relationship between target coherence and performance was stronger after more incongruent trials in accuracy (D) but not RT (E). **F)** Regression estimates for the current target coherence by previous distractor congruence interaction, within each experiment. Error bars on behavior reflect within-participant SEM, error bars on regression coefficients reflect 95% CI. Psychometric functions are jittered on the x-axis for ease of visualization. Feature coherence was rank-ordered and binned into quantiles with equal numbers of trials at each level of target coherence, distractor congruence, or previous distractor congruence.

**Table 2.** Effects of previous conflict on feature sensitivity.

| **Table 2. Effects of previous conflict on feature sensitivity** |  |  |  |  |  |
| --- | --- | --- | --- | --- | --- |
| **DV** | **Predictors** | **Exp 1 (df = 41)**<br>**Effect size (*d*)** | **Exp 2 (df = 15)**<br>**Effect size (*d*)** | **Exp 3 (df = 35)**<br>**Effect size (*d*)** | **Aggregate**<br>***p*-value** |
| **Choice** | Distractor × Prev Distract | 1.59 | 1.45 | 1.74 | **6.15×10<sup>-31</sup>** |
|  | Distractor × Prev Target |  | -0.670 | 0.103 | 0.964 |
|  | Target × Prev Distract |  | -0.473 | -0.990 | **2.83×10<sup>-8</sup>** |
|  | Target × Prev Target |  | 0.418 | 0.644 | **1.25×10<sup>-5</sup>** |
| **Lapse Rate** | Prev Distract | -0.522 | -0.498 | -1.04 | **1.75×10<sup>-10</sup>** |
|  | Prev Target |  | -0.110 | -0.494 | **0.00934** |
| **RT** | Distractor × Prev Distract | -0.836 | -1.44 | -1.79 | **8.99×10<sup>-24</sup>** |
|  | Distractor × Prev Target |  | 0.174 | 0.0618 | 0.520 |
|  | Target × Prev Distract |  | 0.210 | 0.0155 | 0.726 |
|  | Target × Prev Target |  | 0.147 | 0.154 | 0.285 |
|  | Prev Distract | 0.287 | 0.202 | -0.267 | 0.623 |
|  | Prev Target |  | 0.109 | -0.275 | 0.0884 |

Effect sizes are calculated from MAP group-level regression estimates. *P*-values are aggregated across experiments, with statistically significant *p*-values (two-tailed, $\alpha = 0.05$ ) shown in bold.

When varying both target and distractor features (Experiments 2-3), we found an additional influence of previous distractor congruence on target processing, whereby more incongruent previous trials enhanced the influence of target discriminability on the current trial (Figure 3d-e). However, the influence of previous distraction on target processing was substantially smaller than its effect on distractor processing (see Figure 5), and was only found for accuracy (*p* < .001) and not RT (*p* = .57), Finally, we found that performance adapted to the strength of the previous *target*, with less-discriminable targets yielding *lower* sensitivity to target strength (i.e., poorer performance) on the following trial, potentially due to disengagement (Supplementary Figure 2, Table 2). However, like the distractor-target effect, this target-target effect was much smaller than the distractor-distractor effects and only observable in accuracy (*p* < .001) and not RT (*p* = .19).

A common concern when measuring conflict adaptation effects is the extent to which these reflect control adjustment (as typically assumed) or low-level priming that can occur due to stimulus-stimulus or stimulus-response associations (Braem et al., 2019; Hommel et al., 2004; Mayr et al., 2003; Schmidt and De Houwer, 2011). For example, in some tasks, if two adjacent trials are both congruent or both incongruent, they are also more likely to share stimulus-response mappings, biasing analyses of sequential adaptation. Our experiment was designed to largely avoid this potential confound by eliminating stimulus-stimulus repetitions (with two colors assigned to each response hand that never repeat), and by using stochastic motion stimuli (versus, e.g., static arrows) that also have very infrequent exact repetitions (probability of coherence repetition is ∼9%).

However, to further rule out that our key adaptation findings resulted from priming effects, we tested whether adaptation effects were present in our more automatic Attend-Motion blocks. Whereas a priming account would predict similar (within-feature) adaptation effects across both Attend-Color and Attend-Motion blocks, a cognitive control account would predict weaker adaptation effects for Attend-Motion than Attend-Color blocks. We found that adaptation effects during Attend-Motion blocks were overall weak and inconsistently signed (e.g., previous interference led to either increased or decreased sensitivity to distractors across studies; Supplementary Table 4-5). Comparing the adaptation effects across the two types of blocks directly, we found significantly stronger adaptation effects during Attend-Color than Attend-Motion blocks, both for distractors (compared to distractor-distractor adaptation in Attend-Motion; Choice: *p* < .001; RT: *p* < .001) and for motion per se (compared to target-target adaptation in Attend-Motion; Choice: *p* < .001; RT: *p* = .34; Supplementary Table 5). Together, these results suggest that the adaptation effects we observed during Attend-Color trials likely reflected changes in control states rather than stimulus-driven priming.

In addition to affecting sensitivity of choices and RTs to individual features (adaptation effects described above), we found that previous target and distractor information also exerted a small but reliable influence on the likelihood that the participant would respond randomly on the next trial (*lapse rate,* see the Regression Analysis subsection in Methods). Specifically, higher levels of distractor incongruence and lower levels of target discriminability increased subsequent lapse rates (*p*s < .001; Table 2), though these changes were subtle (e.g., post-congruent lapse rates ranged from 0.023% to 0.13% across studies; post-incongruent lapse rates ranged from 0.13% to 0.41% across studies). We did not otherwise find consistent main effects of previous targets and distractors on choice behavior (i.e., in the direction of a particular response) or on RT.

### Performance incentives preferentially enhance target sensitivity

We found that performance on our task adapted to previous distractor-related interference, and that this influence was observed primarily in subsequent processing of the distractor rather than the target. This may reflect a fundamental bias in the control system towards adjusting distractor processing in our task, but it may also reflect a process that is specialized for conflict adaptation. To disentangle these possibilities, we examined how target and distractor processing are influenced by heightened levels of motivation. In Experiment 3 we incorporated an incentive manipulation, with blocks of trials for which participants could either earn a monetary reward for fast and accurate performance, and blocks where performance was not rewarded (Figure 1d).

We found that participants’ accuracy was more sensitive to target discriminability in rewarded blocks than non-rewarded blocks (*d* = 0.61, *p* < .001; Figure 4a, Table 3). This effect of incentives on target sensitivity was specific to choice and not RTs (*d* = -0.10, *p* = 0.47), though participants were overall faster in rewarded blocks (*d* = -0.41, *p* = 0.0045). Participants were also marginally more likely to make lapses response during rewarded blocks (*d* = 0.244, *p* = 0.092). In terms of distractors, we found that in rewarded blocks participants were less sensitive to distractors in RT (*d* = 0.35, *p* = 0.012), albeit with a small effect size, and that incentives did not significantly influence distractor sensitivity in choice (*d* = -0.016, *p* = 0.91).

**Figure 4.**
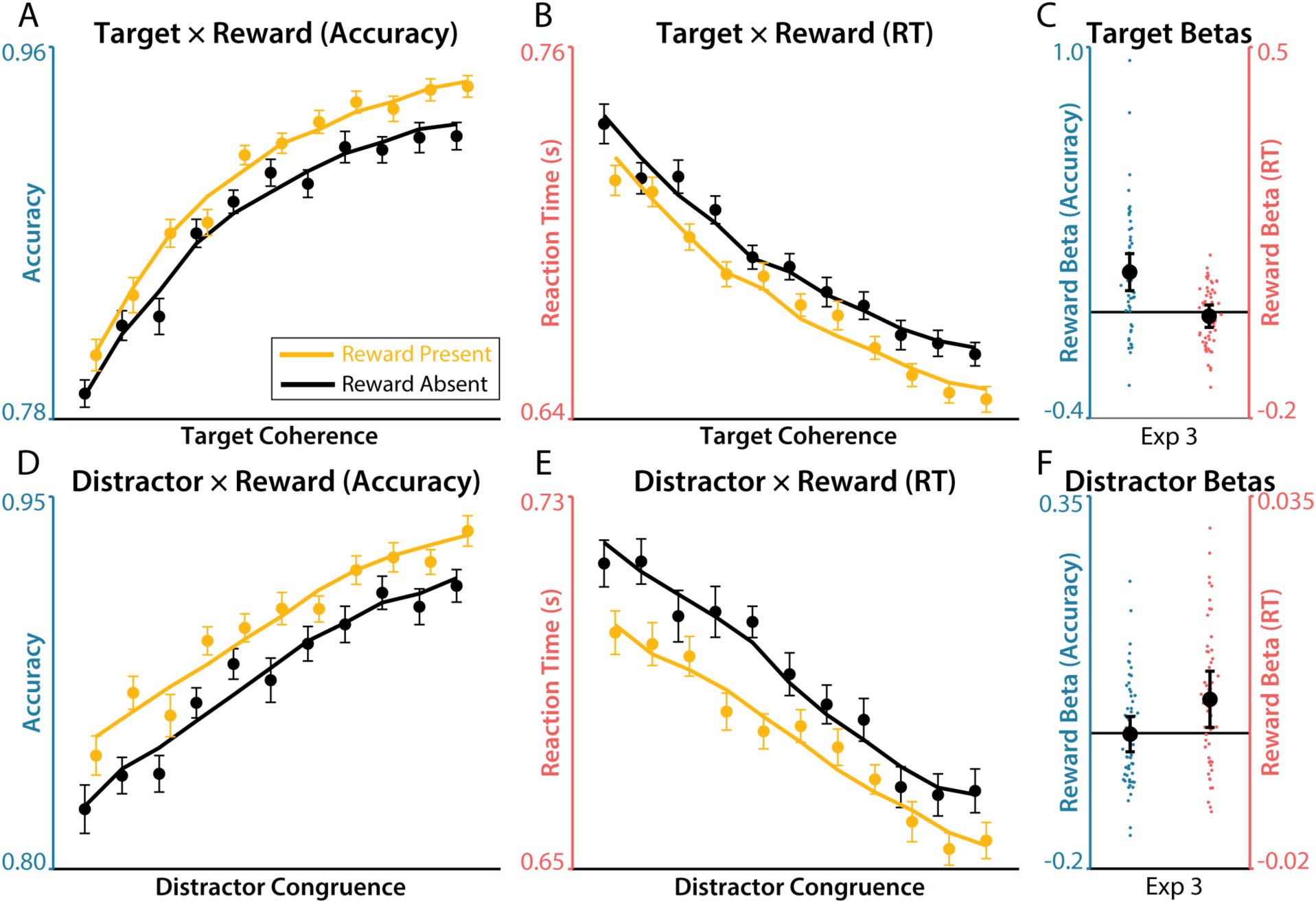
*Influence of incentives on target and distractor sensitivity.* **A-B)** The relationship between target coherence and performance was stronger during incentivized blocks (gold) in the domain of accuracy (A), but not RT (B). Lines depict aggregated regression predictions. **C)** Regression estimates for the target coherence by incentive interaction. **D-E)** The relationship between distractor congruence and performance was weaker on incentivized blocks (gold) in the for RT (E), but not Accuracy (D). **F)** Regression estimates for the distractor congruence by incentive interaction. Error bars on behavior reflect within-participant SEM, error bars on regression coefficients reflect 95% CI. Psychometric functions are jittered on the x-axis for ease of visualization.

**Table 3.** Effects of incentives on feature sensitivity.

| Table 3. Effects of incentives on feature sensitivity |  |  |  |
| --- | --- | --- | --- |
| DV | Predictors | Exp 3 (df = 41)<br>Effect size ($d$) | $p$-value |
| Choice | Target × Reward | 0.612 | $8.56 \times 10^{-5}$ |
| | Distractor × Reward | -0.0156 | 0.911 |
| Lapse Rate | Reward | 0.244 | 0.0924 |
| **RT** | Target × Reward | -0.103 | 0.467 |
| | Distractor × Reward | 0.349 | **0.0195** |
|  | Reward | -0.411 | **0.00447** |

Effect sizes are calculated from MAP group-level regression estimates. Statistically significant $p$ -values (two-tailed, $\alpha = 0.05$ ) are shown in bold.

We further found that the target-enhancing effects of incentives also were not specific to the color dimension. When motion was the target dimension (attend-motion blocks), incentives preferentially increased sensitivity to motion coherence (*d* = 0.70, *p* < .001). Interestingly, incentives had an even larger influence on target sensitivity in attend-motion relative to attend-color blocks (*t*(59.0) = 2.14, *p* = 0.036; Supplementary Table 6-7).

### Previous conflict and incentives have dissociable influences on target and distractor processing

Our within-trial results demonstrated that participants are sensitive to target and distractor information, with little interaction between these dimensions. Consistent with this putative independence, we found that previous interference primarily influenced distractor sensitivity (suppressing distractor sensitivity after trials with incongruent distractors), and that rewards primarily influenced target sensitivity (enhancing target sensitivity when incentivized). These findings strongly suggest a dissociation between target and distractor processing.

To confirm these findings, we formally tested the double dissociation between how incentives and previous interference influenced target and distractor choice sensitivity (Figure 5). We found that previous conflict had a larger absolute effect on distractor processing than it did on target processing in both accuracy (*t*(31.4) = 9.54, *p* = 8.36 × 10^-11^) and RT (*t*(33.7) = 4.64, *p* = 5.14 × 10^-5^). We found that rewards conversely had a larger influence on targets than distractors in Accuracy (*t*(44.5) = 5.08, *p* = 7.22 × 10^-6^), though not in RT (*t*(37.7) = 0.25, *p* = 0.80). The cross- over interaction was also significant in both Accuracy (*t*(39.6) = 10.2, *p* = 1.36 × 10^-12^) and RT (*t*(48.3) = 3.11, *p* = 0.0031), supporting dissociable control over different dimensions of feature processing.

**Figure 5.**
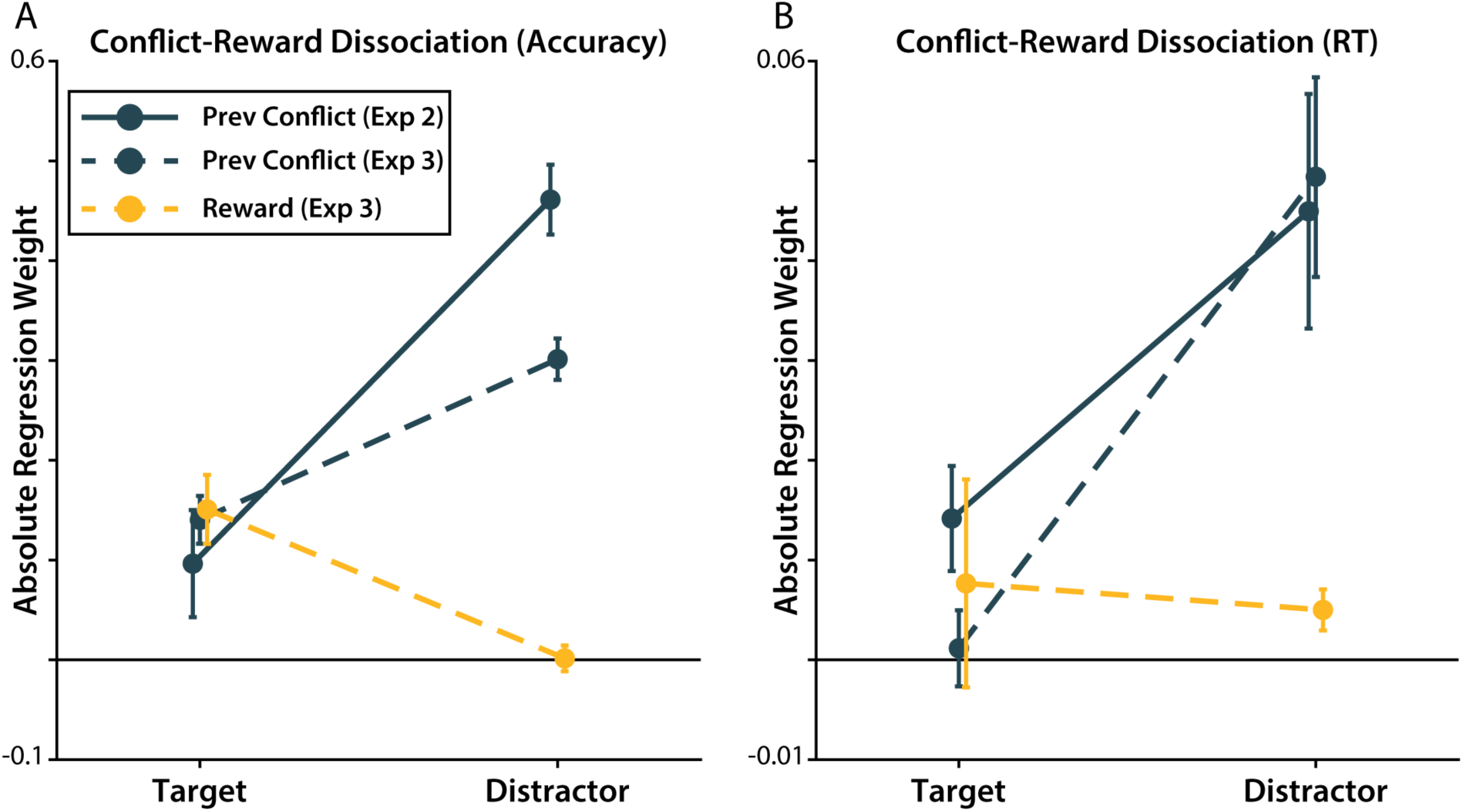
*Dissociations between previous conflict and incentive effects.* Post-conflict effects were significantly larger on distractor sensitivity than target sensitivity in accuracy (A) and RT (B). In contrast, reward effects were significantly larger on target sensitivity than distractor sensitivity in accuracy (A) and similarly large in RT (B). Errors bars show MAP SEM.

These findings are consistent with a pervious neuroimaging experiment that found incentives enhanced responses in target-related areas (visual word form area for text targets) and mostly- incongruent blocks suppressed responses in distractor-related areas (fusiform face area for face distractors; (Soutschek et al., 2015). In the following sections, we extend these convergent findings to explore how previous conflict and incentives influence the dynamics of control implementation.

### Differential within-trial dynamics of target and distractor processing

Our initial results show that participants independently control their sensitivity to target and distractor information. However, previous research has revealed that participants’ task processing also dynamically changes within a trial (Servant et al., 2014; Weichart et al., 2020; White et al., 2011), including in response to incentives (Adkins and Lee, 2021). Whereas much of the previous research has focused on dynamics in spatial attention during flanker tasks (e.g., a shrinking spotlight of attention; (Weichart et al., 2020; White et al., 2011), less is known about the dynamics of attention between features of conjunctive stimuli like those in our task, where target and distractor processing may be more independent (Adkins and Lee, 2021; Servant et al., 2014).

To test how sensitivity to target and distractor features changed within each trial, we measured whether the influence of coherence on participants’ choices depended on reaction time (i.e., the *choice ∼ coherence × RT* interaction). These analyses work under the logic that faster RTs reflect earlier epochs of information processing, consistent with ‘delta function’ analyses of how congruence effects differ across RT quantiles (De Jong et al., 1994; Ridderinkhof, 2002; van den Wildenberg et al., 2010). In the current study, we extended this methodology to study parametric changes in both target and distractor sensitivity over time.

At the earliest RTs, participants were the least sensitive to targets (Figure 6a) and the most sensitive to distractors (Figure 6d). At later RTs, participants became more sensitive to targets (*ds* = 0.69 to 0.97, *p* < .001), and less sensitive to distractors (*ds* = -0.71 to -1.5, *p* < .001; Table 4). This is consistent with an attentional control process that enhances sensitivity to goal-relevant features and suppresses attention towards goal-irrelevant features. Notably, these results suggest that this attentional process occurs ‘online’ within the course of a trial.

**Figure 6.**
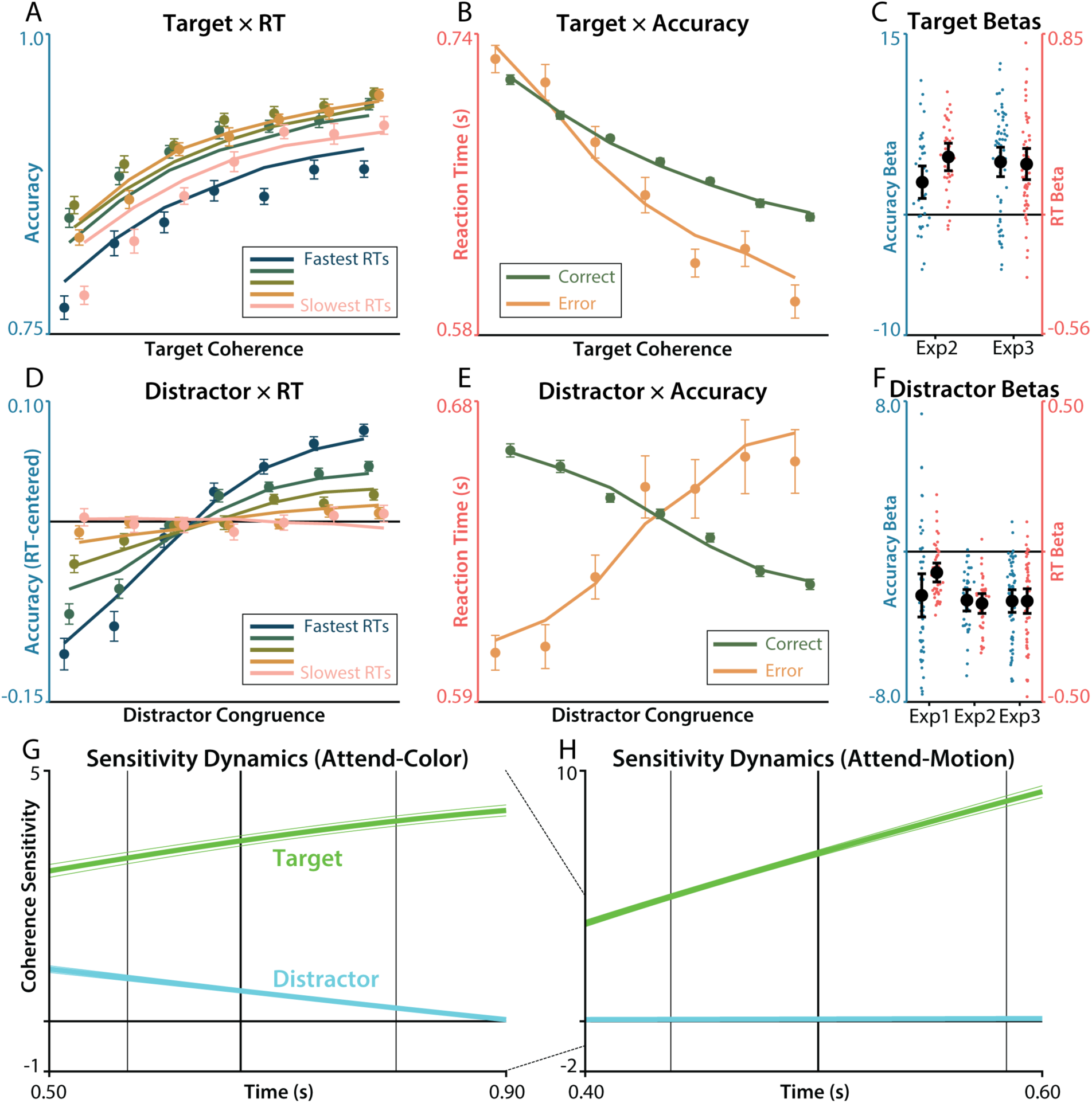
*Target and distractor sensitivity dynamics.* **A)** The relationship between target coherence and accuracy increased at later RTs (pinker color). **B)** Participants responded faster on error trials that correct trial when target coherence was higher. **C)** Regression estimates for the interaction between target coherence and RT (blue) and accuracy (red), within each experiment. **D)** The relationship between distractor congruence and accuracy decreased at later RTs (pinker). Note that these data are mean-centered within each RT bin to remove the target effects in (A) from this visualization of distractor sensitivity. **E)** Participants responded faster on error trials than correct trials when distractors were incongruent. **F)** Regression estimates for the interaction between distractor congruence and RT (blue) and accuracy (red), within each experiment. **G)** Target (green) and distractor (cyan) sensitivity plotted as a function of reaction time, as estimated by our regression model in Attend-Color blocks. Vertical lines indicate quartiles of the RT distribution. **H)** Same as G, but generated from regression models fit to the Attend-Motion blocks. Note the different scaling of the x-axis and y-axis (see dashed line between plots). Error bars on behavior reflect within-participant SEM, error bars on sensitivity estimates reflect between-participant SEM of the predictions, error bars on regression coefficients reflect 95% CI. Psychometric functions are jittered on the x-axis for ease of visualization. Feature coherence and RT were rank-ordered and binned into quantiles with equal numbers of trials at each level of target coherence, distractor congruence, or RT bin.

**Table 4.** Dynamics of feature sensitivity across response times.

| DV | Predictors | Exp 1 (df = 38)<br>Effect size (*d*) | Exp 2 (df = 21)<br>Effect size (*d*) | Exp 3 (df = 41)<br>Effect size (*d*) | Aggregate<br>*p*-value |
| --- | --- | --- | --- | --- | --- |
| **Choice** | Target × RT | | 0.686 | 0.975 | **$4.61 \times 10^{-11}$** |
| | Distractor × RT | -0.709 | -1.54 | -1.19 | **$8.40 \times 10^{-20}$** |
| **Lapse Rate** | RT | 0.247 | 0.928 | 0.534 | **$9.25 \times 10^{-13}$** |
| **RT** | Target × Accuracy | | 1.46 | 0.889 | **$1.99 \times 10^{-14}$** |
| | Distractor × Accuracy | -0.683 | -1.82 | -1.09 | **$1.39 \times 10^{-20}$** |

Effect sizes are calculated from MAP group-level regression estimates. *P*-values are aggregated across experiments, with statistically significant *p*-values (two-tailed, $\alpha = 0.05$ ) shown in bold.

We also fit a complementary analysis for RT (i.e., the *RT ∼ coherence × accuracy* interaction). We found that participants had steeper target coherence slopes on error trials (*ds* = 0.89 to 1.5, *p* < .001; Figure 6b), driven by faster errors when the targets were high coherence, consistent with participants responding before their maximal target sensitivity. Likewise, we found that the relationship between RT and distractor congruence inverted on error trials (*ds* = -0.68 to -1.8, *p* < .001; Figure 6e), with participants making faster errors on more incongruent trials, consistent with an early sensitivity to distractors that is suppressed over time.

These findings suggest online dynamics in the allocation of top-down attention to facilitate target processing and suppress distractor processing, but it is possible that they instead reflect dynamics inherent to the bottom-up processing of color and motion information. To rule out this alternative hypothesis, we tested whether similar sensitivity dynamics were present during Attend-Motion blocks, when color information serves as a much less potent distractor. During these blocks, we found that participants enhanced target (motion) sensitivity faster than they did during Attend-Color blocks (*p* < .001; Figure 6h; Supplementary Table 8-9). In contrast, participants had slower distractor sensitivity dynamics during Attend-Motion blocks (*p* < .001). Together these results demonstrate that these sensitivity dynamics depend on the task that participants are performing, rather than being exclusively due to stimulus-driven factors.

### Previous conflict and incentives influence early trial dynamics

We found that, within a trial, participants dynamically adjusted attention depending on the task at hand, with increasing sensitivity to task-relevant information and decreasing sensitivity to task-irrelevant information over the course of a trial. This raises the question whether the two forms of adaptation we observed, related to previous conflict and incentives, influenced different components of the within-trial attentional dynamics.

To address this question, we first examined how the dynamics of target and distractor sensitivity were altered by the congruence of the distractor on the previous trial (i.e., *Choice ∼ PreviousDistractor × RT × Coherence*). We found that after incongruent trials, participants started the next trial more sensitive to targets and less sensitive to distractors (Figure 7a).

**Figure 7.**
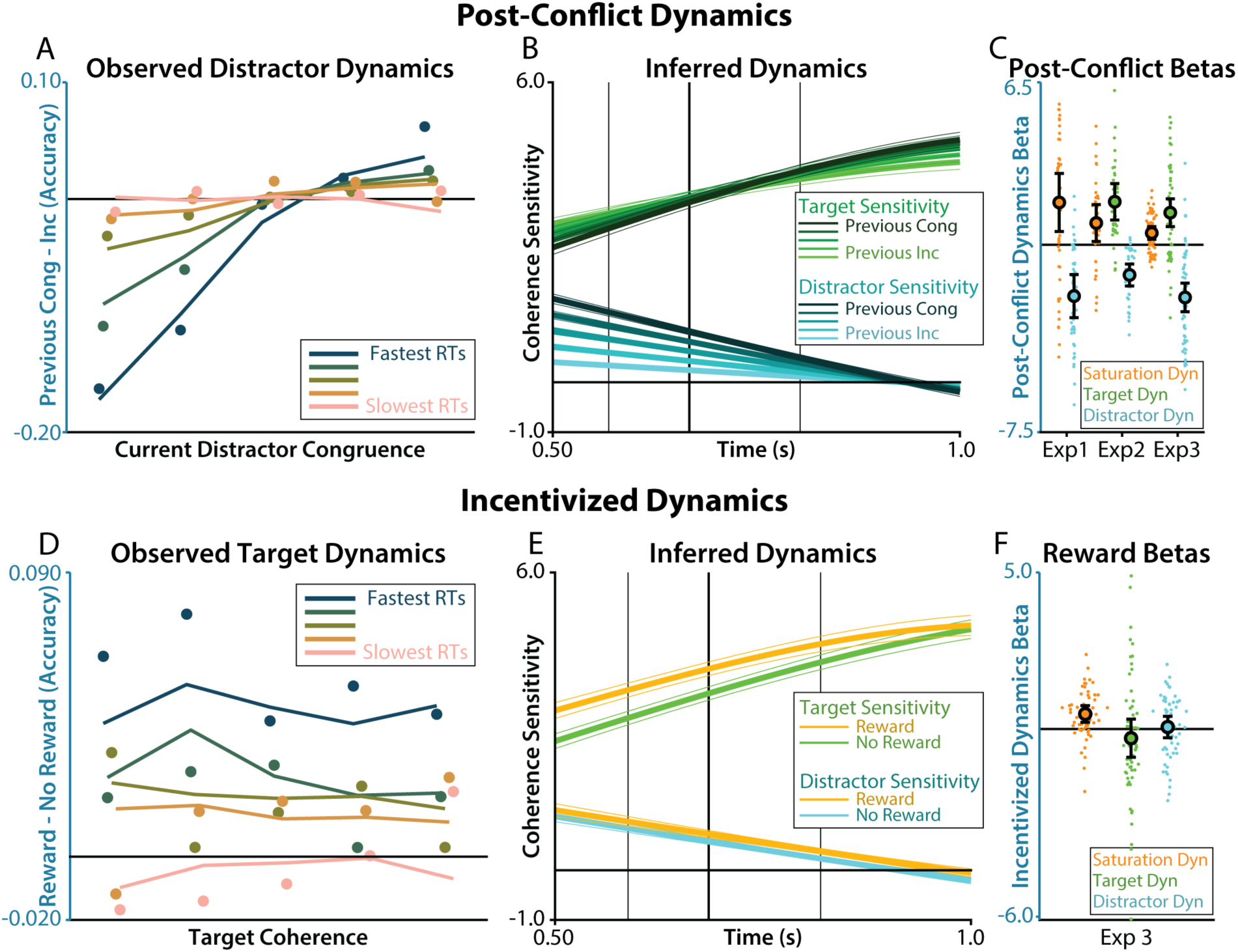
*Influence of conflict and incentives on sensitivity dynamics.* **A)** The relationship between previous distractor congruence and current distractor congruence was strongest for early RTs (bluer color). The y-axis depicts the difference in accuracy between the extreme tertiles of previous congruence, for visualization purposes. **B)** Target (green) and distractor (cyan) sensitivity plotted as a function of previous congruence (color shade) and reaction time (x-axis), as estimated by our regression model. Vertical lines indicate quartiles of the RT distribution. **C)** Regression estimates for the interactions between reaction time and previous congruence on lapse rate (orange); or reaction time, previous congruence, and feature coherence on accuracy (target is green, distractor is cyan). **D)** The relationship between incentives and target coherence was strongest for early RTs (bluer color). The y-axis depicts the difference in accuracy between blocks where there were rewards vs blocks without rewards. **E)** Target (green) and distractor (cyan) sensitivity plotted as a function of incentives (gold) and reaction time (x-axis), as estimated by our regression model. Vertical lines indicate quartiles of the RT distribution. **F)** Regression estimates for the interactions between reaction time and incentives on lapse rate (orange); or reaction time, incentives, and feature coherence on accuracy (target is green, distractor is cyan). Error bars on behavior reflect within-participant SEM, error bars on sensitivity estimates reflect between-participant SEM on the predictions, error bars on regression coefficients reflect 95% CI. Psychometric functions are jittered on the x-axis for ease of visualization. Feature coherence and RT were rank-ordered and binned into quantiles with equal numbers of trials at each level of target coherence, distractor congruence, or RT bin.

Although this means that after congruent trials participants had worse initial conditions (starting less sensitive to targets and more sensitive to distractors), they appeared to compensate for this early disadvantage with faster increases in target enhancement (*ds* = 0.65 to 1.0, *p* < .001) and distractor suppression (*ds* = -0.68 to -1.1, *p* < .001; Table 5). Both post-congruent and post- incongruent trials thus reached similar asymptotic levels of feature sensitivity. This early influence of previous conflict on congruence sensitivity is consistent with previous experiments on the timecourse of conflict adaptation (Stins et al., 2008; Wylie et al., 2010). The current work extends these findings to show both target sensitivity dynamics and conflict-dependent adaptation to target sensitivity dynamics.

**Table 5.**
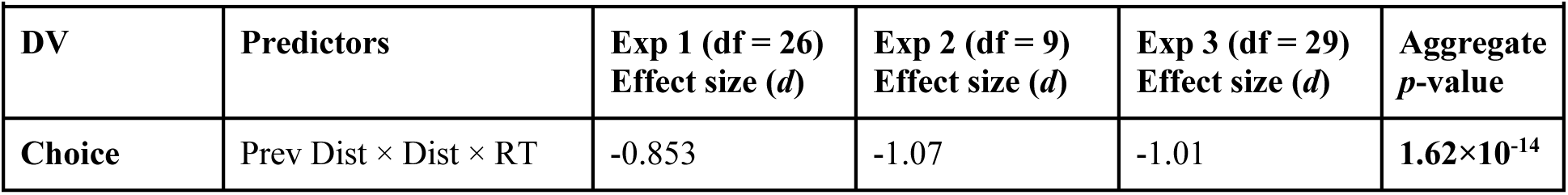
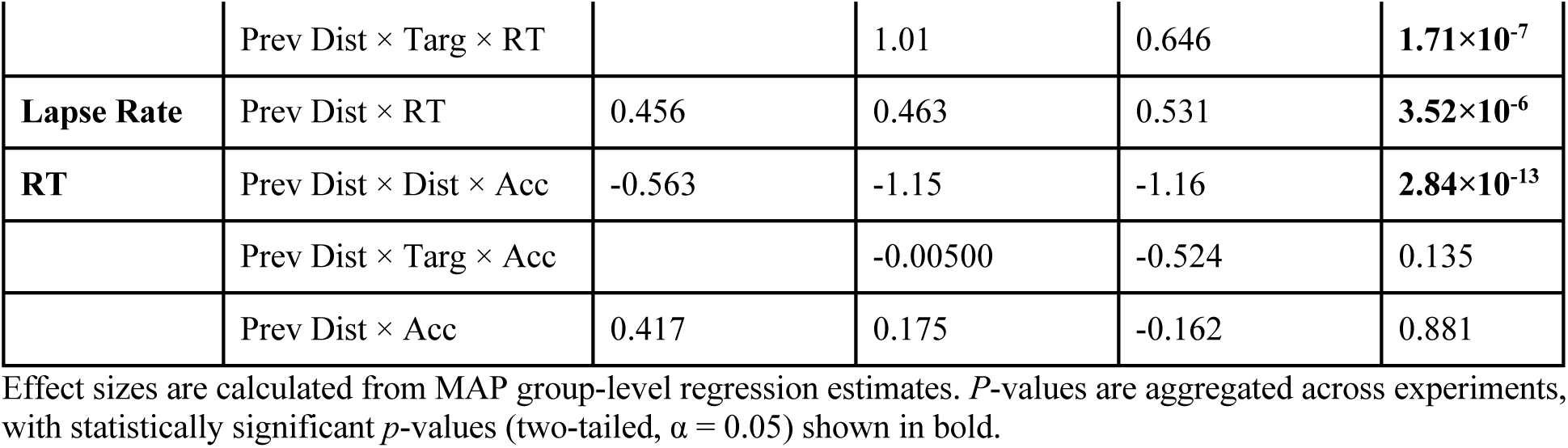
Effects of previous conflict on feature sensitivity dynamics.

We performed the equivalent analysis for incentive-related adaptation (i.e., *Choice ∼ Reward × RT × Coherence*). We found that during incentivized blocks, participants’ initial target sensitivity was higher than during non-incentivized blocks, and remained so across much of the trial (see Figure 7d). However, target sensitivity eventually reached an asymptote, such that towards the end of the trial both incentivized and non-incentivized trials had similar levels of target sensitivity (see slowest quantile in Figure 7d). This convergence was accounted for by larger increases in lapse rates later in incentivized trials (*d* = 0.52, *p* < .001; Table 6). The dynamics of distractor sensitivity, by contrast, did not significantly differ between incentivized and non-incentivized trials (*d* = 0.055, *p* = 0.71).

**Table 6.** Effects of incentives on feature sensitivity dynamics

| Table 6. Effects of incentives on feature sensitivity dynamics |  |  |  |
| --- | --- | --- | --- |
| DV | Predictors | Exp 3 (df = 29)<br>Effect size (*d*) | *p*-value |
| **Choice** | Rew × Target × RT | -0.139 | 0.331 |
| | Rew × Distractor × RT | 0.0546 | 0.712 |
| **Lapse Rate** | Rew × RT | 0.524 | **0.000937** |
| **RT** | Rew × Target × Acc | 0.437 | **0.00432** |
| | Rew × Distractor × Acc | -0.170 | 0.260 |
| | Rew × Acc | -0.540 | **0.000646** |

Effect sizes are calculated from MAP group-level regression estimates. Statistically significant *p*-values (two-tailed, $\alpha = 0.05$ ) are shown in bold.

### An accumulator model of attentional control over target and distractor processing

Our results demonstrate that participants independently control the initialization and online adjustment of attention towards target and distractor features. To parsimoniously account for this set of findings, we developed an accumulator model that integrated elements of previous models used to separately account for performance in tasks involving perceptual discrimination (Gold and Shadlen, 2007; Ratcliff and McKoon, 2008) and overriding prepotent distractors (Cohen et al., 1990; Weichart et al., 2020; White et al., 2011). We used a variant of a feedforward inhibition model, in which inputs provide excitatory inputs to associated response units and inhibitory inputs to alternative response units (Shadlen and Newsome, 2001). Our decision model takes as inputs the color and motion coherence in support of different responses, nonlinearly transforms these inputs, and then integrates evidence for each response in separate rectified accumulators with balanced feedforward excitation and inhibition (Figure 8). The signal-to-noise ratio of the intermediate layer’s outputs are determined by control units that determine the gain of a given feature (Cohen et al., 1990; Musslick et al., 2019). We hand-tuned the parameters of this model to determine whether it could capture our core experimental findings across choice and reaction time.

**Figure 8.**
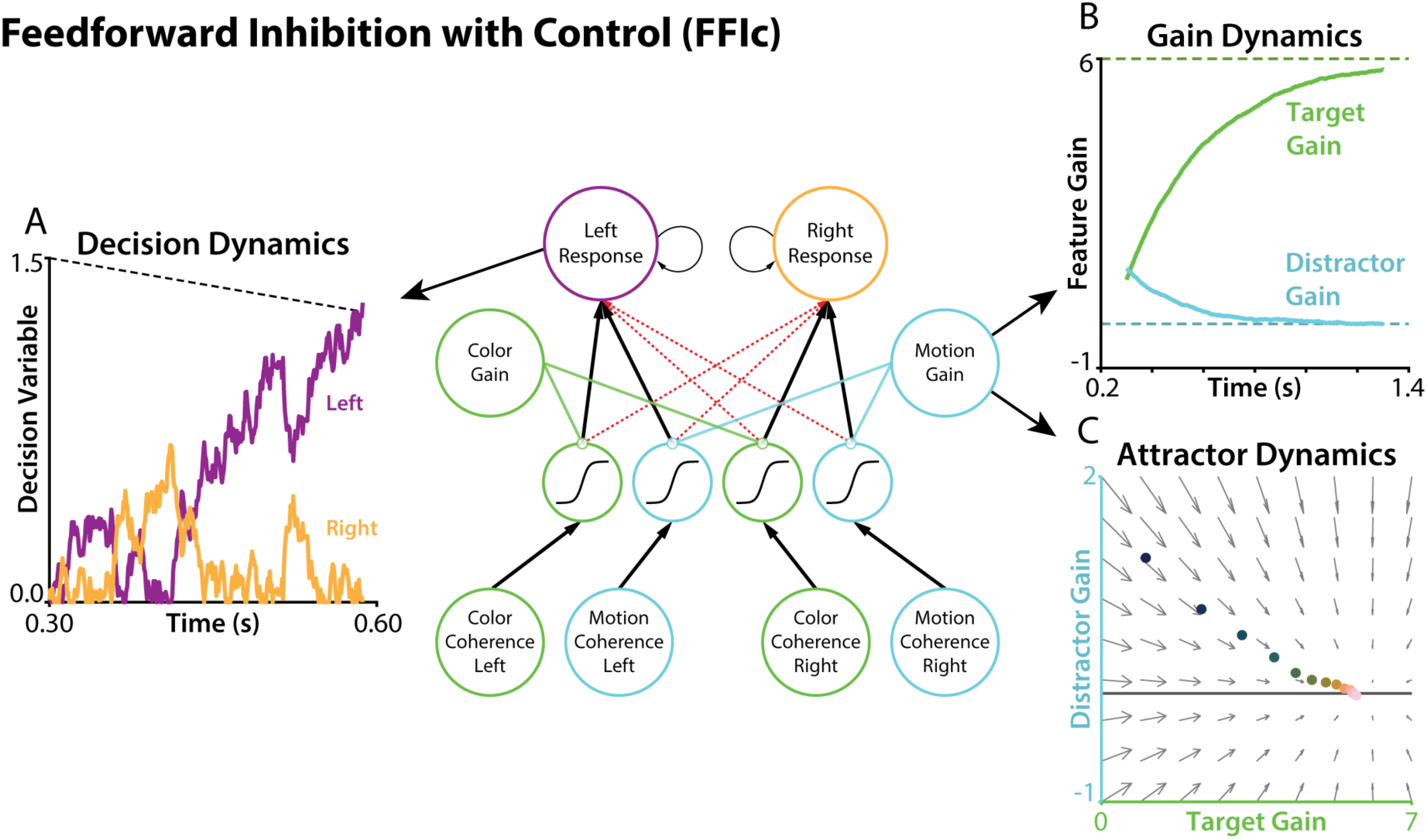
*Feedforward inhibition with control.* Color evidence (green) and motion evidence (blue) are transformed and accumulated to make a choice. Balanced excitatory connections (black solid lines) and inhibitory connections (red dashed lines) cause accumulation of the difference in evidence for each response. **A)** Evidence for the left response (purple) and right response (orange) are accumulated over time without leak. When one of the accumulators crosses a (linearly collapsing) decision threshold, the model chooses that response. **B)** Within each trial, the signal-to-noise of each feature pathways is controlled by a feature gain. Over time within a trial, the feature gains for targets (green) and distractors (cyan) exponentially approach to a fixed level (high gain for targets, zero gain for distractors). Note the difference in x-axis scaling compare to Figure 6G. **C)** An equivalent visualization of the dynamics in B. Attractor dynamics drive target and distractor gains to their fixed level, shown at different timepoints within the trial (pinker colors are later in the trial). The horizontal line depicts zero distractor gain.

Our accumulator model was able to reproduce our key within-trial findings. During our main Attend-Color trials, it generated responses that were faster and more accurate with increasing color coherence (Figure 9a) and slower and less accurate with increasing motion incongruence (Figure 9b). We simulated Attend-Motion trials by increasing the target gain and decreasing the distractor gain, to capture potential differences in both automaticity and control. Now, our model generated responses that were even faster and more accurate with increasing target coherence (now motion; Figure 9c) but that were insensitive to distractor congruence (now color; Figure 9d), replicating the main behavioral results in Attend-Motion blocks. Notably, distractor effects were not reproduced in an accumulator competition model parameterized to be more ‘race-like’ (Supplementary Figure 3; (Teodorescu and Usher, 2013). This occurred because larger inputs (whether congruent or incongruent) drove faster reaction times.

**Figure 9.**
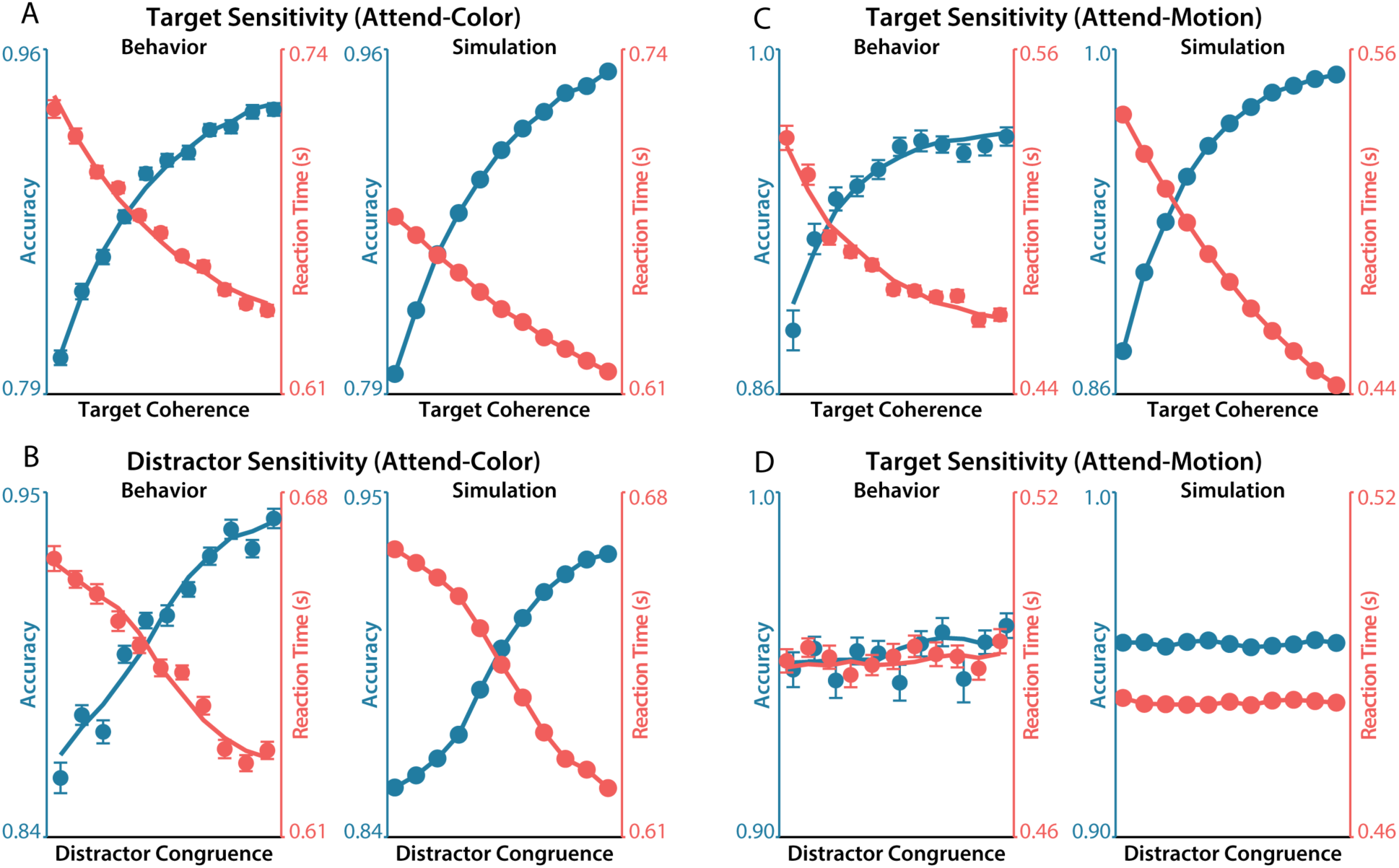
*Simulation of target and distractor sensitivity (see Figure 2).* **A-B)** Sensitivity to target coherence (A) and distractor congruence (B) in behavior (left) and in the FFIc simulation (right) for Attend-Color blocks. **C-D)** Same as A-B, but for Attend-Motion blocks.

We next used this model to test potential mechanisms underlying participants’ within- and between-trial control adaptations. First, we tested whether participants’ apparent within-trial dynamics in feature sensitivity plausibly resulted from actual within-trial changes in control gains governing feature sensitivity, or whether such dynamics could result from static control gains. We implemented time-varying feature gains as attractors with an initial gain (e.g., reflecting bottom-up salience or learning) that exponentially approaches a fixed point (e.g., determined by the task goals and control; cf. (Musslick et al., 2019; Steyvers et al., 2019). We found that incorporating these time-varying gains into our accumulator model allowed it to reproduce participants’ behavioral dynamics. In accuracy, our model replicated the shift in target sensitivity over time, with the collapsing bound reducing performance on the slowest trials (Figure 10a). Our model similarly captured participants’ decreased target sensitivity at later RTs (Figure 10b). Finally, our model recreated the analogous effects in RT, with faster errors for high coherence targets and incongruent distractors (Figure 10c-d). Critically, we were unable to replicate these qualitative patterns of behavior with FFI models in which control gains that were frozen throughout the trial (Supplementary Figure 4); drift diffusion models with across-trial variability in gain, noise, or threshold; or drift diffusion models with within-trial dynamics in noise or threshold (Supplementary Figure 5).

**Figure 10.**
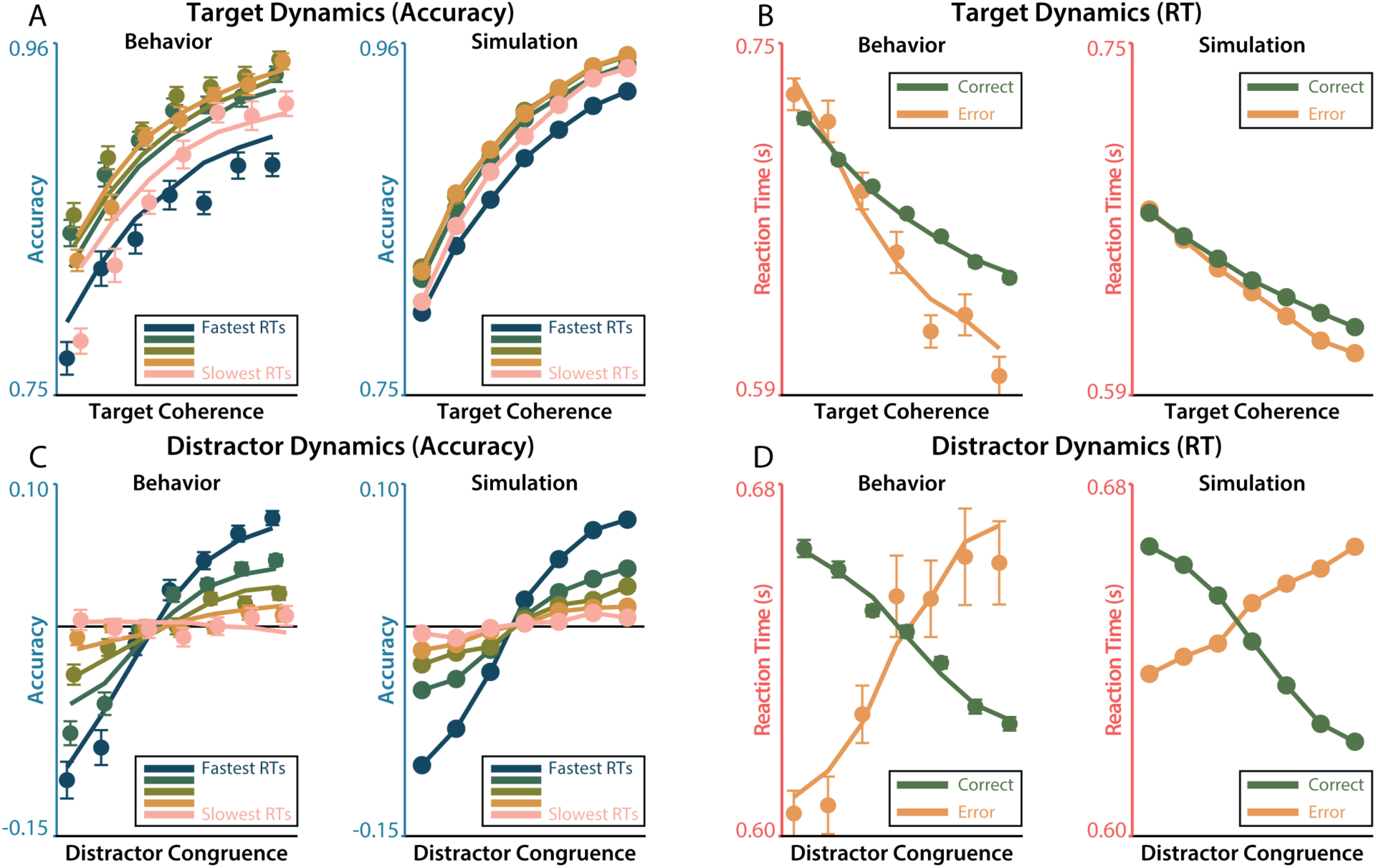
*Simulation of target and distractor sensitivity dynamics (see Figure 6).* **A-B)** RT-dependent (A) and accuracy-dependent (B) sensitivity to target coherence in behavior (left) and in the FFIc simulation (right). **C-D)** Same as A-B, but for distractor congruence.

At later RTs, participants were more likely to exhibit lapses in performance (i.e., choose randomly; ds = 0.25 to 0.93, *p* < .001, see Table 4), which were estimated with a separate term in our regression models (see the Regression Analyses subsection in Methods). This is evident in a ‘hook’ that emerges at the slowest RT bin, whereby trials in this bin demonstrate lower overall accuracy than those in adjacent RT bins (see Figure 10a, left panel, pink line). This hook is often observed in RT-conditioned accuracy, with gradually better performance followed by poorer performance for the slowest RTs (van den Wildenberg et al., 2010; Weichart et al., 2020). Our simulation captured this global reduction in accuracy by including a collapsing boundary (Drugowitsch et al., 2012; Rosenbaum et al., 2022), which leads to late errors irrespective of feature coherence (see Supplementary Figure 5 for contrast to fixed bound). Notably, even though overall accuracy is reduced over time, target sensitivity is stronger at the slowest RT bin relative to the earliest RT bin (compare navy and pink psychometric slopes in Figure 10a), consistent with both feature-selective and global dynamics.

The parallel feature pathways in this model are designed to capture the independent influences of a target and distractor information (Lindsay and Jacoby, 1994). However, the time-varying feature gains providing an account for the weak interactions we observed between target and distractor sensitivity in accuracy. Despite there being no competition in feature processing in our model, we found these weak target-distractor interactions emerge in *simulated* accuracies but not simulated RTs. This interaction appeared to result from the different time courses of target and distractor sensitivity. As in participants’ behavior, the model’s errors due to incongruent distractors tend to occur early (Figure 10c-d), censoring target processing at a lower (early) level of sensitivity. This interplay between feature sensitivity *dynamics* (but not overall feature sensitivity per se) offers a plausible explanation for the subtle and seeming inconsistent interactions in participants’ behavior.

Having provided an account of how each of our stimulus features is processed over the course of the trial depending on the task goal, we next tested a potential model-based account of the two forms of control adaptation we observed across trials. Our participants demonstrated enhanced target sensitivity on rewarded blocks, and suppressed distractor sensitivity after increasingly incongruent trials. In both cases, adaptation appeared to enhance sensitivity to stimulus features at the fastest reaction times. To account for these findings, we modified the initial conditions of our model’s gain dynamics (Figure 11a). We simulated post-interference adaptation by initializing the distractor gain closer to its asymptote, and we simulated reward incentivization by initializing the target gain closer to its asymptote. We found that these simulations qualitatively reproduced participants’ behavior, with stronger adaptation and reward effects earlier in the trial than later. The exponential dynamics in our attractor network parsimoniously accounts for the fact that dynamics tended to be faster when they were initialized further from the fixed point (i.e., post-congruent trials). Thus, our model was able to capture the range of findings in this experiment: target-distractor sensitivity, within-trial dynamics, and how the dynamics of target and distractor processing may be influenced by control.

**Figure 11.**
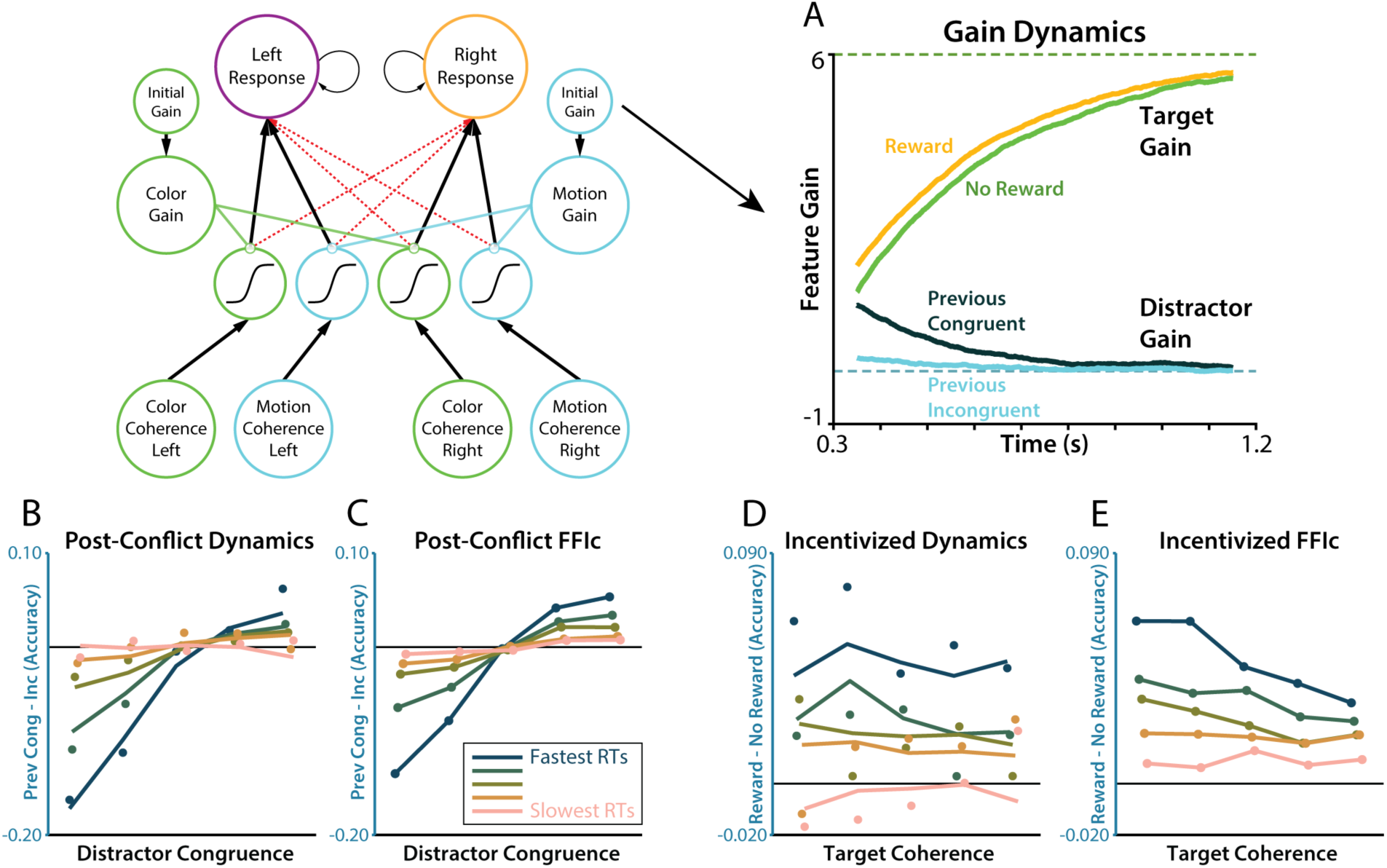
*Simulation of post-conflict and incentive effects (see Figure 7).* **A)** The influences of previous congruence (shade) and incentive effects (gold) were implemented through changes to the initial conditions of the feature gain dynamics, with previous congruence influencing initial distractor gain and incentives influence initial target gain. **B-C)** The influence of previous congruence on distractor sensitivity dynamics in behavior (B) and in the FFIc simulation (C). **D-E)** The influence of incentives on target sensitivity dynamics in behavior (D) and in the FFIc simulation (E).

## Discussion

When faced with distraction, we can sustain good performance by engaging with relevant information or ignoring disruptive information. Our experiment revealed that these strategies are under independent cognitive control, and are driven by distinct attentional dynamics. Using a bivalent random dot motion task with parametric target and distractor coherence (PACT), we found that target and distractor information have independent influences on participants’ performance. Furthermore, we found that participants’ sensitivity to targets and distractors was preferentially modulated by incentives and previous interference, respectively. These adaptations altered the initial conditions of feature-selective gains, which was followed by dynamic enhancement to target gains and suppression of distractor gains. These behavioral phenomena could be parsimoniously explained by a hybrid sequential sampling model with goal-dependent attractor dynamics over feature weights.

Together, these results support a cognitive control architecture that is parametric, multivariate, and dynamic. Previous research has found that cognitive effort is enhanced in response to incentives (Parro et al., 2018; Yee and Braver, 2018) and to previous conflict (Egner, 2007; Gratton et al., 1992). The current experiments extend these previous findings to show that these adaptations are both graded in their intensity, and selective in their allocation. These findings are consistent with a multivariate perspective on cognitive control (Ritz et al., 2022), in which people optimize a configuration of control signal according to their costs and benefits (Musslick et al., 2015; Shenhav et al., 2013). The target and distractor configurations observed here add to a body of work teasing apart the conditions under which people coordinate across multiple control signals (Danielmeier et al., 2011; Leng et al., 2021; Noonan et al., 2016; Simen et al., 2009; Soutschek et al., 2015; Wöstmann et al., 2019).

A core question arising from these results is why there are preferential relationships between previous conflict with distractors, and incentives with targets. One possibility is that this is due to credit assignment. A system that could properly assign credit to features based on their contribution to conflict and incentives should allocate control towards distractors and targets. Distractors are a salient source of response conflict, and participants could adjust sensitivity to reduce this conflict. When participants were performing the more automatic Attend-Motion blocks, during which response conflict was absent, this adaptation was also absent. In contrast, reward contingencies were explicitly tied to target discrimination performance. During Attend-Motion blocks, there was a stronger association between target coherence and performance (e.g., due to response compatibility, and that only targets contributed to accuracy), potentially explaining why these blocks had larger incentive effects. This account is consistent with Bayesian models of cognitive control, such as those that predict feature congruence (Jiang et al., 2014; Yu et al., 2009) or the value of control policies (Bustamante et al., 2021; Lieder et al., 2018).

Our results also provide insight into the dynamic implementation of attentional control. Previous work has shown that within-trial attentional dynamics play an important role in both decision making (Callaway et al., 2021; Krajbich et al., 2010; Li and Ma, 2021; Westbrook et al., 2020) and cognitive control (Adkins and Lee, 2021; Hardwick et al., 2019; Servant et al., 2014; Weichart et al., 2020; White et al., 2011). These foundational experiments have largely focused on spatial attention, with far less known about the dynamics of feature-based attention, where processing of targets and distractors is less mutually constrained. Furthermore, relatively few experiments have studied how attentional dynamics are modified in response to control drivers like incentives or task demands (Adkins and Lee, 2021; van den Wildenberg et al., 2010; Yu et al., 2009).

Our experiments show that the dynamics of target and distractor sensitivity are independent, and that previous conflict and incentives appear to operate through changes to the initial conditions of these feature gains, rather than their speed or asymptotic level. These findings are broadly consistent with influential theories of attentional dynamics which propose that early task processing is largely driven by feature salience and statistical or reinforcement learning, whereas attentional control has a relatively slower timecourse (Awh et al., 2012; Theeuwes, 2018, 2010, see also van den Wildenberg et al., 2010). If participants are learning the relevance of different features, it’s possible that these initial conditions in part reflect the prior probability that attention towards targets or distractors will support task goals (Lieder et al., 2018; Yu et al., 2009). Similar to how response priors are reflected in the initial decision state (Bogacz et al., 2006; Simen et al., 2009), priors on feature priority may be reflected in the initial attentional state. In the case of previous interference, this could reflect learning whether distractors enhance performance (e.g., after trials on which congruent distractors led to better performance), or a local estimate of the probability a trial will be congruent (Yu et al., 2009). For incentives, this may reflect the expected target-reward contingency. Future research should investigate this account by measuring attentional dynamics as participants learn task contingencies (Shenhav et al., 2018).

Our patterns of conflict- and incentive-dependent dynamics help rule out stimulus-driven dynamics and support independent control over feature processing. After congruent trials, participants started the next trial with more similar target and distractor gains, that were then more quickly separated within the trial (Figure 7b). If these dynamics were an artifact of the decision process (e.g., due to accumulator attractors; (Wong and Wang, 2006), then we would expect that when target and distractor gains are initially more similar, there would be slower dynamics. Instead, we found faster dynamics, supporting a role for feedback control that reconfigures attentional gain to align with task goals. Additionally, during incentivized blocks, we saw that participants modified attentional dynamics for targets, but not distractors. This finding further supports the independence of these attentional dynamics, demonstrating that participants can alter attention towards individual features one at a time. This pattern of incentives enhancing sensitivity to target information, while also producing faster responding and a marginally higher lapse rate, is consistent with previous work on motivated attention. A previous experiment used drift diffusion modeling to show that participants increase their rate of evidence accumulation and decrease their response threshold when faced with higher rewards, consistent with the reward-rate optimal policy (Leng et al., 2021). The current experiment extends these findings by revealing how specific attentional adjustments improve evidence accumulation, providing a more process-oriented account of motivated cognitive control.

Our dynamical process model may help link behavior in response conflict tasks to cognitive dynamics in other domains. In the domain of task-switching, recent cognitive models have developed similar dynamical accounts of how people reconfigure task sets. Classic work has shown that switch costs exponentially decay with preparation time (Monsell and Mizon, 2006; Rogers and Monsell, 1995), similar to the dynamics in the current task. Computational models have formalized these task sets dynamics during the switch preparation period (Musslick et al., 2019; Ueltzhöffer et al., 2015) and across trials (Steyvers et al., 2019). If the within-trial dynamics we observe reflect a “task set micro-adjustment” (Ridderinkhof, 2002), then this modeling approach may help unify switching dynamics across executive domains. Interestingly, control over initial conditions also plays a central role in the neural dynamics of motor preparation (Churchland et al., 2010; Remington et al., 2018b, 2018a), highlighting the similarities across motor and cognitive domains (Ritz et al., 2022, 2020) and offering potential neural predictions for our task.

The evidence we provide for dissociable control over target and distractor processing is consistent with previous neuroscience experiments that used neural correlates of stimulus processing to argue for independent enhancement and suppression processes (Gazzaley et al., 2005; Noonan et al., 2016; Soutschek et al., 2015; Wöstmann et al., 2019). Our results extend these findings by exploring how different factors can contribute to dynamic reconfiguration of target and distractor attention, which we formalize in an explicit process model. Notably, our findings diverge from neuroimaging experiments that have suggested that control primarily acts through enhancements to target processing (Egner and Hirsch, 2005). One potential source of this divergence may be that people’s control strategies differ depending on the source of task conflict (Braem et al., 2014; Egner, 2008; Egner et al., 2007). For example, tasks evoking stimulus-stimulus conflict (e.g., semantic competition in Stroop task) may require different strategies than tasks evoking stimulus-response conflict (e.g., distractors driving competing responses, as in PACT). Although previous work using Stoop-like tasks has found similar patterns of control adjustments as in the current experiment (Soutschek et al., 2015), this raises the broader question of how the effects we found may generalize outside the current experiment. According to the Expected Value of Control theory, and the Learned Value of Control model that builds upon it, control strategies are adapted to specific task contexts (Lieder et al., 2018; Ritz et al., 2022; Shenhav et al., 2013). The current results show that participants can independently control target and distractor processing when these features are independent, and future work should explore whether control strategies appropriately accommodate other kinds of tasks.

Our analyses of attentional dynamics depend on participants’ own response times and choices, raising concerns about selection biases. While evidence accumulation modelling typically depends on choice-conditioned reaction times, inferring time-varying influence of targets and distractors presents a particular challenge. To address these concerns, we used simulations to show that the dynamic profiles we observed cannot be accounted for by an evidence accumulation model with static gains on target and distractor processing (Supplementary Figure 4) or models with dynamic changes to non-selective components like decision threshold (Supplementary Figure 5). Introducing dynamic feature gains allowed us to account for those same patterns (Figures 9 to 11; Supplementary Figure 5). These results are consistent with previous work validating DDM estimates of attentional dynamics in conflict tasks (White et al., 2018, 2011). Even if these measurements are valid, using sparse behavioral measures is an inefficient method for measuring latent dynamics, and may combine multiple processes (e.g., accumulation and threshold adjustments). By integrating across multiple convergent measures of decision and attentional dynamics – including interrogation protocols (Adkins and Lee, 2021; Hardwick et al., 2019), motor tracking (Erb et al., 2016; Menceloglu et al., 2021; Scherbaum et al., 2010), and/or temporally-resolved neuroimaging (Fischer et al., 2018; Scherbaum et al., 2011; Weichart et al., 2020; Yeung et al., 2004) – future work can help strengthen and build on our understanding of continuous changes in the configuration of multiple control processes.

The evidence accumulation modeling in the current experiment was able to categorically rule out several alternative architectures, demonstrating the necessity and sufficiency of feature-specific adjustments for capturing the full array of putative attentional dynamics. Our model validation approach supports our interpretation of feature-selective adjustments, while committing less strongly to the specific formulation of attentional control (e.g., a specific model parameterization, or the functional form of the collapsing bound). An important direction for future research should be to leverage emerging methods for parameter estimation to directly fit our accumulator model to participants’ behavior (Fengler et al., 2021; Weichart et al., 2020).

This approach will help extend insights from the current experiment, such as enabling participant-specific parameters to reveal individual differences in attentional control.

This experiment provides new insight into how we flexibly adapt to the changing demands of our environment. We find that this flexible control aligns information processing with task goals, and can be captured by a well-defined process model. The developments from this experiment can help extend models of cognitive control towards richer accounts of how multivariate control configurations, such as across targets and distractors, are optimized during goal-directed behavior.

## Methods

### Participants

All participants provided informed consent in compliance with Brown University’s Institutional Review Board, participating for either course credit or pay. We excluded participants from our analyses if they had <70% accuracy during attend-color blocks or completed less than half of the experiment. Fifty-seven individuals participated in Experiment 1 (mean(SD) age: 20.6(2.21); 36 female; 1 excluded), 42 individuals participated in Experiment 2 (age: 19.1(0.971); 31 female; 2 excluded), and 62 individuals participated in Experiment 3 (age: 19.8(1.38); 47 female; 2 excluded), resulting in 156 included participants across the three experiments. Sample sizes were guided by piloting in Experiment 1 and experimental standards in cognitive control research (commonly n = 20-40; e.g., Adkins and Lee, 2021; Danielmeier et al., 2011; Jiang et al., 2015; Vogel et al., 2020; White et al., 2011).

### Parametric Attentional Control Task (PACT)

We developed the Parametric Attentional Control Task (PACT), extending tasks used to study decision-making (Kang et al., 2021; Mante et al., 2013; Shenhav et al., 2018) and cognitive control (Danielmeier et al., 2011). On each trial, participants viewed an array of moving dots (i.e., random dot kinematogram), presented in one of four colors (see Figure 1). Participants were taught to match two colors to a left keypress and two colors to a right keypress (with colors counterbalanced across participants). The majority color did not repeat on adjacent trials to avoid priming (Braem et al., 2019).

The direction of the dot motion (leftward or rightward) was task-irrelevant and could be consistent with the color response (distractor congruent trials) or it could be inconsistent with this response (distractor incongruent trials). Uniquely in this experiment, we parametrically varied the degree of distractor congruence on each trial by varying the motion coherence (percentage of dots moving in the same direction). Distractor congruence was linearly spaced between 95% congruence and 95% incongruence, sampled randomly across trials. We made the motion highly salient to maximize the conflict induced by this distracting dimension (Wöstmann et al., 2021), with dots moving quickly across a large aperture. Critically, our primary interest is understanding how people adjust sensitivity to salient distractors, and not the main effect of distractor congruence per se. (p. 27)

In Experiment 1, all of the dots were the same color (100% color coherence), creating a parametric extension of the Simon conflict tasks (Danielmeier et al., 2011). In Experiments 2 and 3, the dots contained a mixture of two colors associated with different responses. Color coherence was linearly spaced between 65% to 95%, drawn randomly across trials.

To maintain the salience of the motion dimension throughout the session (Shiffrin and Schneider, 1977), participants alternated between blocks of the task above (‘attend-color’ trials, putatively more control-demanding) and blocks where participants were instructed to instead indicate the direction of the dot motion (‘attend-motion’ trials, putatively less control-demanding). Mirroring the attend-color blocks, in Experiment 1 we held the motion coherence constant (maximal) during attend-motion blocks, while varying the color coherence across trials. In Experiments 2 and 3, we varied the coherence of both dimensions during attend-motion blocks.

Comparing performance across tasks that are matched for visual and motoric demands also allows us to test whether behavioral effects depend on stimulus or response confounds. For example, participants’ behavior may be influenced by eye movement confounds (e.g., bottom-up attentional capture from to motion coherence), response repetition biases (e.g., due to responses switching more often than repeating), or stimulus-response priming (e.g., due to how response switching coincides with stimulus transitions). Critically, Attend-Color and Attend-Motion tasks differ in their putative control demands, allowing us to isolate stimulus-response confounds from goal-directed control.

### Session

Participants first performed 100 motion-only training trials (0% coherent color) and 100 color-only training trials (0% coherent motion; order counterbalanced across participants) to learn the stimulus-response mappings. During training, participants received accuracy feedback on every trial. During the main experiment, participants performed two types of interleaved blocks, without trial-wise feedback. Participants alternated between longer attend-color blocks (100 trials) and shorter attend-motion blocks (Experiment 1: 20-50 trials; Experiment 2-3: 30 trials; order counterbalanced across participants). In Experiments 1 and 2, at the end of each block participants were told their average accuracy and median RT, and encouraged to respond quickly and accurately. Participants were not given this information in Experiment 3 to avoid interactions with the incentive manipulation (see below). Participants took self-timed breaks between blocks.

### Stimuli

Participants were seated approximately 60cm from a computer screen, making their responses on a customizable gaming keyboard in a dark testing booth. The random dot motion array was presented in the center of the screen (∼15 visual degrees in diameter; ∼66.8 dots per visual degree squared; 19” LCD display at 60Hz). The dots colors were approximately (uncalibrated) isoluminant and perceptually equidistant (RGB: [187, 165, 222], [150, 180, 198], [192, 169, 168], [157, 184, 130]; (Teufel and Wehrhahn, 2000) and moved at ∼15 visual degrees per second. Each trial started with a random inter-trial interval (Experiment 1: 0.5 – 1.5s; Experiment 2-3: 0.5 – 1.0s). There was an alerting cue 300ms before the trial onset, indicated by the fixation cross turning from grey to white, to minimize non-decision time. The stimuli were then presented until either a response was made, or a deadline was reached (Experiment 1: 3s; Experiment 2-3: 5s).

### Task Variants

Experiment 1: These data incorporate several similar versions of this task developed during piloting. The main differences across versions were the number of distractor congruence levels (mean(range) = 13.5(11-15)), the number of trials per attend-motion block (mean(range) = 26(20-50)), and the total number of trials (mean(range) = 469(300-700) attend-color trials). We did not find significant differences in performance across versions, and so our analyses collapsed across these versions. Experiment 1 also included a learning condition in a separate set of blocks, which was outside the scope of the current paper and not included in the analyses we report.

Experiments 2 & 3: These data come from a single task variant (though see Experiment 3’s incentive manipulation below). In this variant, we presented participants with 11 levels of target coherence and 11 levels of distractor congruence, linearly spaced within their coherence range and randomly sampled across trials. Participants performed 12 blocks of 100 attend-color trials interleaved with 12 blocks of 30 attend-motion trials.

### Incentivized Variant (Experiment 3)

In Experiment 3, we informed participants before the main session that they would be able to earn a monetary reward for good performance. On ‘Reward’ blocks, we would randomly select trials, and if they were both fast and accurate on those trials, they would receive a bonus payment. On ‘No Reward’ blocks, participants would not be eligible to earn a reward. We indicated the incentive condition at the beginning of each block with a label and text coloring (gold text for ‘Reward’, white text for ‘No Reward’). Participants performed Attend-Color and Attend-Motion blocks in one incentive condition before alternating to the other incentive condition (order counterbalanced across participants). At the end of the experiment, participants received a bonus calculated from their performance (mean(SD) bonus: $2.5($0.57)USD).

### Regression Analyses

We used a hierarchical nonlinear regression of choice and reaction time as a tractable and minimally theory-laden measure of performance (Supplementary Figure 1). We designed these regression models to quantify changes to target and distractor sensitivity, while controlling for global factors like behavioral autocorrection and how task factors may change lapse rates. The results of these regression analyses then provided the basis for our explicit process modeling (see below). We confirmed that our regression models are identifiable using Belsley collinearity diagnostics (collintest in MATLAB; Supplementary Table 10).

In particular, we implemented hierarchical expectation maximization (EM) in MATLAB R2020a (using emfit; available at github.com/mpc-ucl/emfit) to provide a maximum a posteriori (MAP) estimates for the mean and covariance of parameters linking task features to performance. This fitting algorithm alternates between finding the MAP estimates of participants’ parameters given the current group-level expectations (M-step; with 5 parameter re-initialization per step), and updating this group-level expectation based on participants’ estimated parameters (E-step), repeated until convergence. We fit separate regression to each experiment for independent replications of our findings. Analysis code is available at github.com/shenhavlab/PACT-public.

Our regression approach simultaneously estimated parameters for choice and RT:

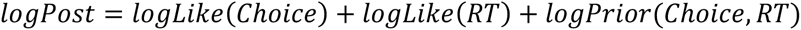

Our choice sub-function used a lapse-logistic likelihood function:

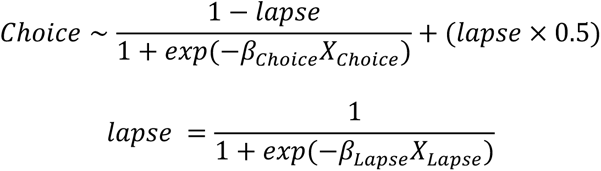

Where *β_Choice_* and *β_Lapse_* are parameter vectors, and *X_Choice_* and *X_Lapse_* are design matrices. Our RT sub-function used a shifted lognormal likelihood function:

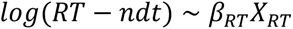

Where again *β_RT_X_RT_* is a linear model, and *ndt* is the estimated non-decision time. Rare RTs less than *ndt* were assigned a small likelihood.

Finally, the prior probability of the parameters was evaluated under a multivariate normal distribution defined by the group-level parameter mean and covariance, improving the robustness of our estimates through regularization. Critically, we estimated this group-level covariance across both choice and RT parameters, which better regularized our estimates and produced a joint model of performance at the group level.

All regression design matrices included an intercept (choice bias or average RT), an autoregressive component (previous trial’s choice or RT), and the transformed target and distractor coherence (scaled between -1 and 1). We transformed feature coherences using a saturating nonlinearity,

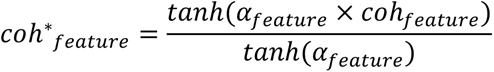

with *α_target_* and *α_distracor_* fit as free parameters. This approach distinguishes the coherence nonlinearity (*α*) from how strongly coherence influences performance (*β_Choice_* and *β_RT_*), with our analyses focused on the latter. To constrain these *α* parameters, we estimated one parameter for both choice and RT, capturing similar non-linearities across both performance measures.

In our more complex models (e.g., incentives), our primary focus was on how additional task features moderated the influence of tanh-transformed feature coherence on performance. Lower order effects of moderating factors (e.g., previous distractor congruence) were included in the lapse rate for choice analysis, and as a main effect in RT analyses. The full parameter sets for all analyses are available in Supplementary Data.

We excluded trials in our regression if they were 1) the first trial of the block, 2) shorter than 200ms or longer than 2s, 3) occurred after an error or after a trial was too fast/slow and 4) in reaction time analyses, if the current trial was an error. These exclusion criteria were chosen to be inclusive, while avoiding trials where there were likely to be a mixture of different cognitive processes (e.g., post-error adjustments).

We performed statistical inference on the parameters using an estimate of the group-level error variance from the emfit package, necessary to avoid violations of independence across participants from our hierarchical modelling. Contrast tests across models used Welsh’s (unequal variance) t-tests, with contrasts weighting studies by the square root of the sample size. We aggregated *p*-values across studies using Lipták’s method (Lipták, 1958; Zaykin, 2011), weighting studies by the square root of their sample size. Correlations between parameters were calculated by converting the group-level MAP covariance matrix to a correlation matrix.

We generated posterior predictive checks (trend lines on figures) by generating model predictions for all trials, and then aggregating these predictions in the same way as participants’ behavior. This approach allows us to distinguish whether our model systematically deviates from behavior from whether deviations are driven by variability in parameters across participants. To provide finer-grained insights into our model fit, we generated additional posterior predictive checks that aggregate trends across all participants (Supplementary Figure 6) and that highlight single participants (Supplementary Figure 7).

We generated sensitivity dynamics plots (e.g., Figure 6) by computing the regression-estimated coherence effect conditioned on RT. For a range of simulated RTs, the estimated motion sensitivity timeseries is:

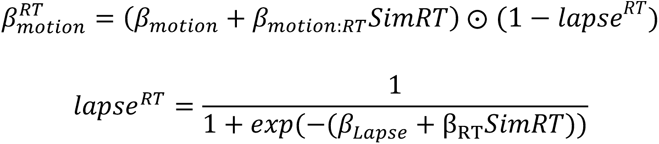

Where *βs* are regression weights estimated in our analysis, *SimRT* is a vector of simulated RTs (e.g., .5:.01:1), and ⊙ indicates element-wise multiplication. For control-dependent dynamics (i.e., incentivized dynamics; see Figure 7), we included 2-way and 3-way interactions between feature sensitivity, RT, and control drivers. We generated these sensitivity dynamics for each participant, and then plotted the mean and between-participant standard error.

### Feedforward Inhibition with Control Model

To provide a bridge between our regression analyses and processes models of decision-making, we adopted a generative modeling approach and tested whether participants behavior could be reproduced by a sequential sampling model (Figure 8). See simulation code for full parameter set at github.com/shenhavlab/PACT-public.

This model takes as inputs the color and motion coherence in support of different responses (e.g., *coh_colorleft_*), nonlinearly transforms these inputs (e.g., *coh*_colorleft_*; see regression analyses above), and then integrates evidence for each response in separate rectified accumulators (*x_left_* and *x_right_*).

For example, evidence for the left response would be calculated as:

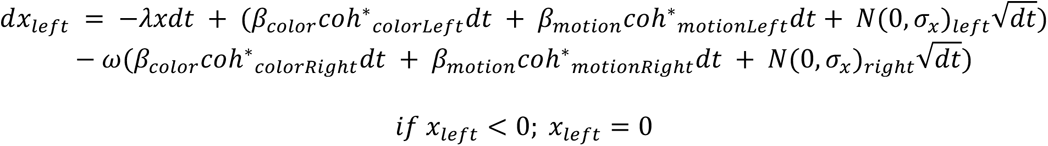

The model makes a choice when one of the accumulators reaches a linearly collapsing decision bound rectified above 0.01. We used a balanced feedforward inhibition model without leak (*λ* = 0 and *ω* = 1), approximating a (rectified) drift diffusion process (Bogacz et al., 2006). Note that parameterizations of a leaky competing accumulator could also approximate the DDM (Bogacz et al., 2007, 2006), and so are plausible alternatives to our implementation. We preferred the FFI model because it provides a simple interpolation between DDM and race-like decision processes.

To capture dynamics in participants’ feature sensitivity, we modified our accumulation model to incorporate an attractor network for the feature weights, a model we call the feedforward inhibition with control model (FFIc model). This control process has a gain setpoint (e.g., aims for zero gain on distractors), and noisily drives the gain from its initial condition (*β*^0^) towards this attractor according to a gain *K*. For example, the motion gain would be governed by:

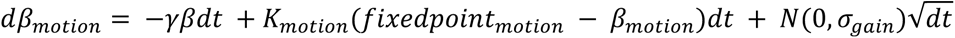

With the leak term γ fixed to 0 as in the decision process.

We simulated 10,000 trials for each combination of target discriminability and distractor congruence (11 x 11 x 10,000), and then aggregated simulated behavior in the same way we aggregated participants’ behavior.

## Supporting information

withinTrial models

priorConflict models

incentives models

## Supplementary Tables

**Supplementary Table 1.** Target and distractor sensitivity (Attend-Motion)

|  |  |  |  |  |  |
| --- | --- | --- | --- | --- | --- |
| DV | Predictors | Exp 1 (df = 45)<br>Effect size (*d*) | Exp 2 (df = 25)<br>Effect size (*d*) | Exp 3 (df = 45)<br>Effect size (*d*) | Aggregate<br>*p*-value |
| Choice | Target coherence |  | 4.39 | 5.44 | **5.69×10<sup>-53</sup>** |
|  | Distractor congruence | 0.732 | 0.352 | 0.269 | **0.000916** |
|  | Target × Distractor |  | 0.0522 | 0.145 | 0.6210 |
| RT | Target coherence |  | -1.47 | -1.63 | **2.71×10<sup>-21</sup>** |
|  | Distractor congruence | 0.125 | 0.179 | 0.407 | 0.121 |
|  | Target × Distractor |  | 0.0710 | 0.0371 | 0.864 |

Effect sizes are calculated from MAP group-level regression estimates. *P*-values are aggregated across experiments, with statistically significant *p*-values (two-tailed, $\alpha = 0.05$ ) shown in bold.

**Supplementary Table 2.**
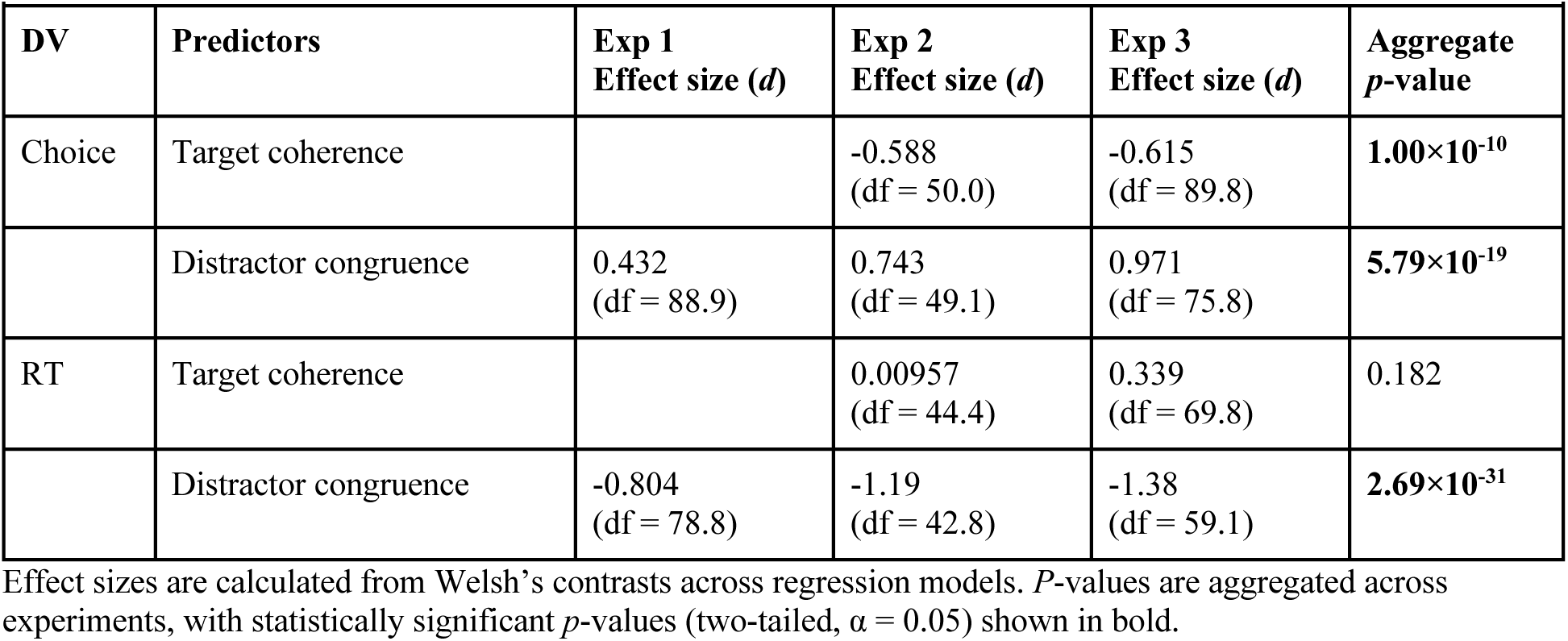
Target and distractor sensitivity (Attend-Color - Attend-Motion)

|  |  |  |  |  |  |
| --- | --- | --- | --- | --- | --- |
| DV | Predictors | Exp 1<br>Effect size (*d*) | Exp 2<br>Effect size (*d*) | Exp 3<br>Effect size (*d*) | Aggregate<br>*p*-value |
| Choice | Target coherence |  | -0.588<br>(df = 50.0) | -0.615<br>(df = 89.8) | **1.00×10<sup>-10</sup>** |
|  | Distractor congruence | 0.432<br>(df = 88.9) | 0.743<br>(df = 49.1) | 0.971<br>(df = 75.8) | **5.79×10<sup>-19</sup>** |
| RT | Target coherence |  | 0.00957<br>(df = 44.4) | 0.339<br>(df = 69.8) | 0.182 |
|  | Distractor congruence | -0.804<br>(df = 78.8) | -1.19<br>(df = 42.8) | -1.38<br>(df = 59.1) | **2.69×10<sup>-31</sup>** |

Effect sizes are calculated from Welsh's contrasts across regression models. *P*-values are aggregated across experiments, with statistically significant *p*-values (two-tailed, $\alpha = 0.05$ ) shown in bold.

**Supplementary Table 3.**
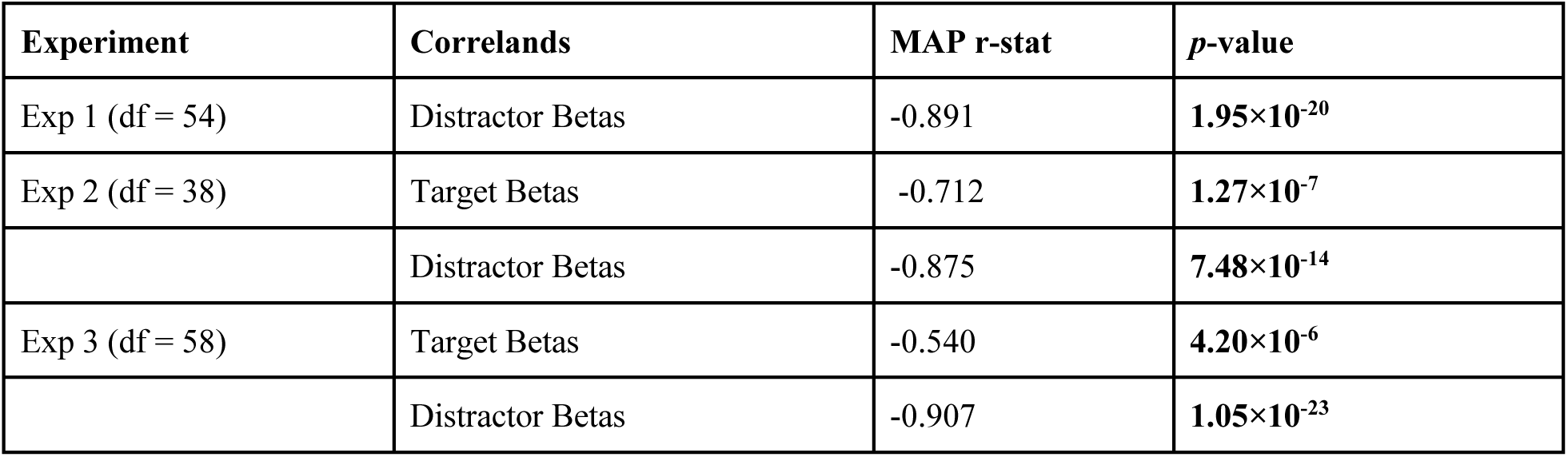
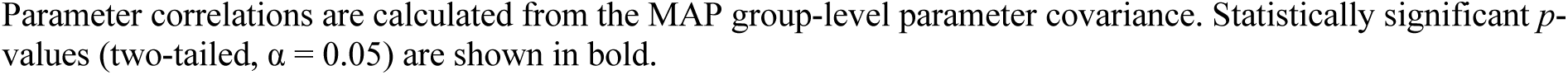
Correlations between RT and accuracy betas

| Supplementary Table 3. Correlations between RT and accuracy betas |  |  |  |
| --- | --- | --- | --- |
| Experiment | Correlands | MAP r-stat | *p*-value |
| Exp 1 (df = 54) | Distractor Betas | -0.891 | **1.95×10<sup>-20</sup>** |
| Exp 2 (df = 38) | Target Betas | -0.712 | **1.27×10<sup>-7</sup>** |
|  | Distractor Betas | -0.875 | **7.48×10<sup>-14</sup>** |
| Exp 3 (df = 58) | Target Betas | -0.540 | **4.20×10<sup>-6</sup>** |
|  | Distractor Betas | -0.907 | **1.05×10<sup>-23</sup>** |

Parameter correlations are calculated from the MAP group-level parameter covariance. Statistically significant $p$ -values (two-tailed, $\alpha = 0.05$ ) are shown in bold.

**Supplementary Table 4.** Effects of previous conflict on feature sensitivity (Attend-Motion)

|  |  |  |  |  |  |
| --- | --- | --- | --- | --- | --- |
| DV | Predictors | Exp 1 (df = 41)<br>Effect size ($d$) | Exp 2 (df = 19)<br>Effect size ($d$) | Exp 3 (df = 39)<br>Effect size ($d$) | Aggregate<br>$p$-value |
| Choice | Distractor × Prev Distract | 0.0113 | -0.176 | 0.293 | 0.578 |
| | target × Prev Target | | -0.00795 | -0.424 | 0.268 |
| RT | Distractor × Prev Distract | 0.503 | -0.427 | 0.181 | 0.291 |
| | target × Prev Target | | 0.131 | -0.0333 | 0.777 |

Effect sizes are calculated from MAP group-level regression estimates. $P$ -values are aggregated across experiments, with statistically significant $p$ -values (two-tailed, $\alpha = 0.05$ ) shown in bold.

**Supplementary Table 5.** Effects of previous conflict on feature sensitivity (Attend-Color - Attend-Motion)

|  |  |  |  |  |  |
| --- | --- | --- | --- | --- | --- |
| DV | Predictors | Exp 1<br>Effect size ($d$) | Exp 2<br>Effect size ($d$) | Exp 3<br>Effect size ($d$) | Aggregate<br>$p$-value |
| Choice | Distractor-dependent<br>(Distractor - Distractor) | 0.505<br>(df = 52.7) | 0.984<br>(df = 33.6) | 0.280<br>(df = 47.7) | **$4.39 \times 10^{-8}$** |
|  | Motion-dependent<br>(Distractor - Target) |  | 0.448<br>(df = 21.9) | 0.651<br>(df = 40.2) | **$2.27 \times 10^{-5}$** |
| RT | Distractor-dependent<br>(Distractor - distractor) | -0.629<br>(df = 80.2) | -0.956<br>(df = 29.3) | -1.20<br>(df = 70.7) | **$3.40 \times 10^{-23}$** |
|  | Motion-dependent<br>(Distractor - Target) |  | -0.237<br>(df = 19.1) | -0.0929<br>(df = 39.3) | 0.337 |

Effect sizes are calculated from Welch's contrasts across regression models. $P$ -values are aggregated across experiments, with statistically significant $p$ -values (two-tailed, $\alpha = 0.05$ ) shown in bold.

**Supplementary Table 6.**
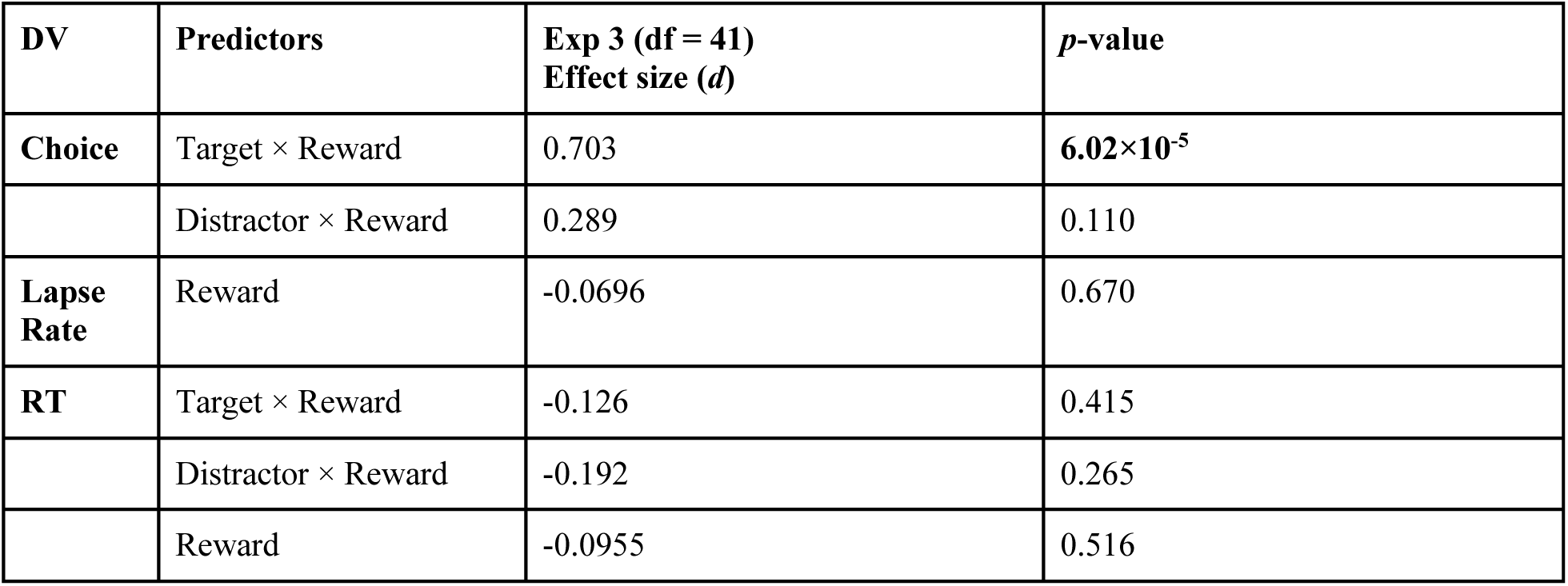
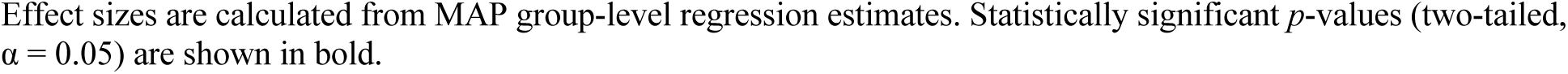
Effects of incentives on feature sensitivity (Attend-Motion)

|  |  |  |  |
| --- | --- | --- | --- |
| DV | Predictors | Exp 3 (df = 41)<br>Effect size ($d$) | $p$-value |
| Choice | Target × Reward | 0.703 | **$6.02 \times 10^{-5}$** |
| | Distractor × Reward | 0.289 | 0.110 |
| Lapse Rate | Reward | -0.0696 | 0.670 |
| RT | Target × Reward | -0.126 | 0.415 |
| | Distractor × Reward | -0.192 | 0.265 |
|  | Reward | -0.0955 | 0.516 |

Effect sizes are calculated from MAP group-level regression estimates. Statistically significant $p$ -values (two-tailed, $\alpha = 0.05$ ) are shown in bold.

**Supplementary Table 7.** Effects of incentives on feature sensitivity (Attend-Color – Attend-Motion)

|  |  |  |  |
| --- | --- | --- | --- |
| DV | Predictors | Exp 3<br>Effect size ($d$) | $p$-value |
| Choice | Target × Reward | -0.279<br>($df = 59.0$) | **0.0363** |
| | Distractor × Reward | -0.230<br>($df = 48.3$) | 0.117 |
| RT | Target × Reward | 0.0566<br>($df = 58.3$) | 0.667 |
| | Distractor × Reward | 0.271<br>($df = 77.0$) | **0.0199** |

Effect sizes are calculated from Welsh's contrasts across regression models. Statistically significant $p$ -values (two-tailed, $\alpha = 0.05$ ) are shown in bold.

**Supplementary Table 8.** Dynamics of feature sensitivity across response times (Attend-Motion)

|  |  |  |  |  |  |
| --- | --- | --- | --- | --- | --- |
| DV | Predictors | Exp 1 ($df = 38$)<br>Effect size ($d$) | Exp 2 ($df = 21$)<br>Effect size ($d$) | Exp 3 ($df = 41$)<br>Effect size ($d$) | Aggregate<br>$p$-value |
| Choice | Target × RT | | 2.03 | 1.31 | **$3.30 \times 10^{-20}$** |
| | Distractor × RT | -0.257 | 0.194 | -0.490 | **0.0226** |
| Lapse Rate | RT | 0.509 | 0.763 | -0.234 | 0.941 |
| RT | Target × Accuracy | | 0.772 | 0.679 | **$1.59 \times 10^{-7}$** |
| | Distractor × Accuracy | 0.0758 | -0.211 | 0.0199 | 0.931 |

Effect sizes are calculated from MAP group-level regression estimates. $P$ -values are aggregated across experiments, with statistically significant $p$ -values (two-tailed, $\alpha = 0.05$ ) shown in bold.

**Supplementary Table 9.** Dynamics of feature sensitivity across response times (Attend-Color – Attend- Motion)

|  |  |  |  |  |  |
| --- | --- | --- | --- | --- | --- |
| DV | Predictors | Exp 1<br>Effect size ($d$) | Exp 2<br>Effect size ($d$) | Exp 3<br>Effect size ($d$) | Aggregate<br>$p$-value |
| Choice | Target × RT | | -2.08<br>($df = 23.0$) | -1.09<br>($df = 46.1$) | **$3.35 \times 10^{-17}$** |
| | Distractor × RT | -0.102<br>($df = 56.3$) | -0.585<br>($df = 22.4$) | -0.539<br>($df = 78.5$) | **$4.57 \times 10^{-5}$** |
| RT | Target × Accuracy | | -0.429<br>($df = 23.0$) | -0.422<br>($df = 46.2$) | **0.00148** |
|  | Distractor × Accuracy | -0.344<br>(df = 66.4) | -0.648<br>(df = 29.6) | -0.723<br>(df = 78.6) | **4.56×10<sup>-11</sup>** |

Effect sizes are calculated from Welsh's contrasts across regression models. *P*-values are aggregated across experiments, with statistically significant *p*-values (two-tailed, $\alpha = 0.05$ ) shown in bold.

**Supplementary Table 10.** Model Collinearity

| **Supplementary Table 10. Model Collinearity** |  |  |  |
| --- | --- | --- | --- |
| **Model** | **Experiment** | **Accuracy Model Collinearity**<br>**median [25% - 75%]** | **RT Model Collinearity**<br>**median [25% - 75%]** |
| Baseline | Experiment 1 | 1.4 [1.4 – 1.5] | 1.1 [1.1 – 1.1] |
|  | Experiment 2 | 1.4 [1.4 – 1.5] | 1.1 [1.1 – 1.1] |
|  | Experiment 3 | 1.4 [1.4 – 1.5] | 1.1 [1.1 – 1.1] |
| Post-Conflict | Experiment 1 | 1.5 [1.4 – 1.5] | 1.2 [1.1 – 1.3] |
|  | Experiment 2 | 1.4 [1.4 – 1.5] | 1.3 [1.2 – 1.3] |
|  | Experiment 3 | 1.5 [1.4 – 1.5] | 1.3 [1.2 – 1.3] |
| Reward | Experiment 3 | 1.4 [1.4 – 1.5] | 1.2 [1.1 – 1.2] |
| Dynamics | Experiment 1 | 1.5 [1.5 – 1.6] | 1.5 [1.3 – 2.0] |
|  | Experiment 2 | 1.5 [1.4 – 1.5] | 1.6 [1.4 – 1.7] |
|  | Experiment 3 | 1.5 [1.4 – 1.5] | 1.7 [1.5 – 2.0] |
| Post-Conflict Dynamics | Experiment 1 | 1.6 [1.6 – 1.8] | 2.2 [1.7 – 3.3] |
|  | Experiment 2 | 1.5 [1.4 – 1.5] | 2.1 [1.8 – 2.4] |
|  | Experiment 3 | 1.5 [1.5 – 1.6] | 2.1 [1.7 – 2.5] |
| Reward Dynamics | Experiment 3 | 1.5 [1.5 – 1.6] | 2.0 [1.7 – 2.4] |

Belsley collinearity diagnostics for core models (from MATLAB's collintest). Diagnostic values are the ratio of the design matrix's largest singular value to its smallest singular value, summarized at different participant quantiles (i.e., median is the participant at the 50<sup>th</sup> percentile). A value of 1 is perfect orthogonality, and values below 30 are within the default tolerance. All values are well below 30, indicating tolerable collinearity.

## Supplementary Figures

**Supplementary Figure 1. (supplement to Figure 1).**
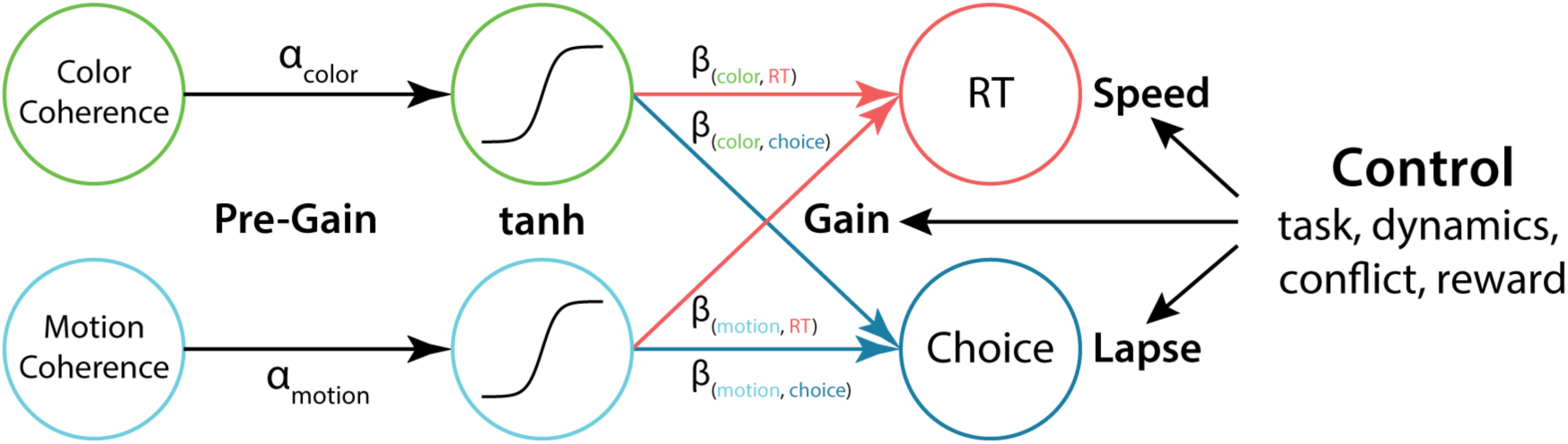
*Regression schematic.* To estimate feature sensitivity, trial-specific color (green) and motion (cyan) coherence levels were passed through a hyperbolic tangent non-linearity (tanh), with the *α* parameter determining the strength of the non-linearity (see Methods). The linear relationships between transformed coherence and performance (RT in red and Choice in blue) were our estimates of participants’ feature sensitivity. Our critical analyses tested whether potential indices of control (e.g., task instructions or incentives) moderated this feature sensitivity.

**Supplementary Figure 2 (supplement to Figure 3).**
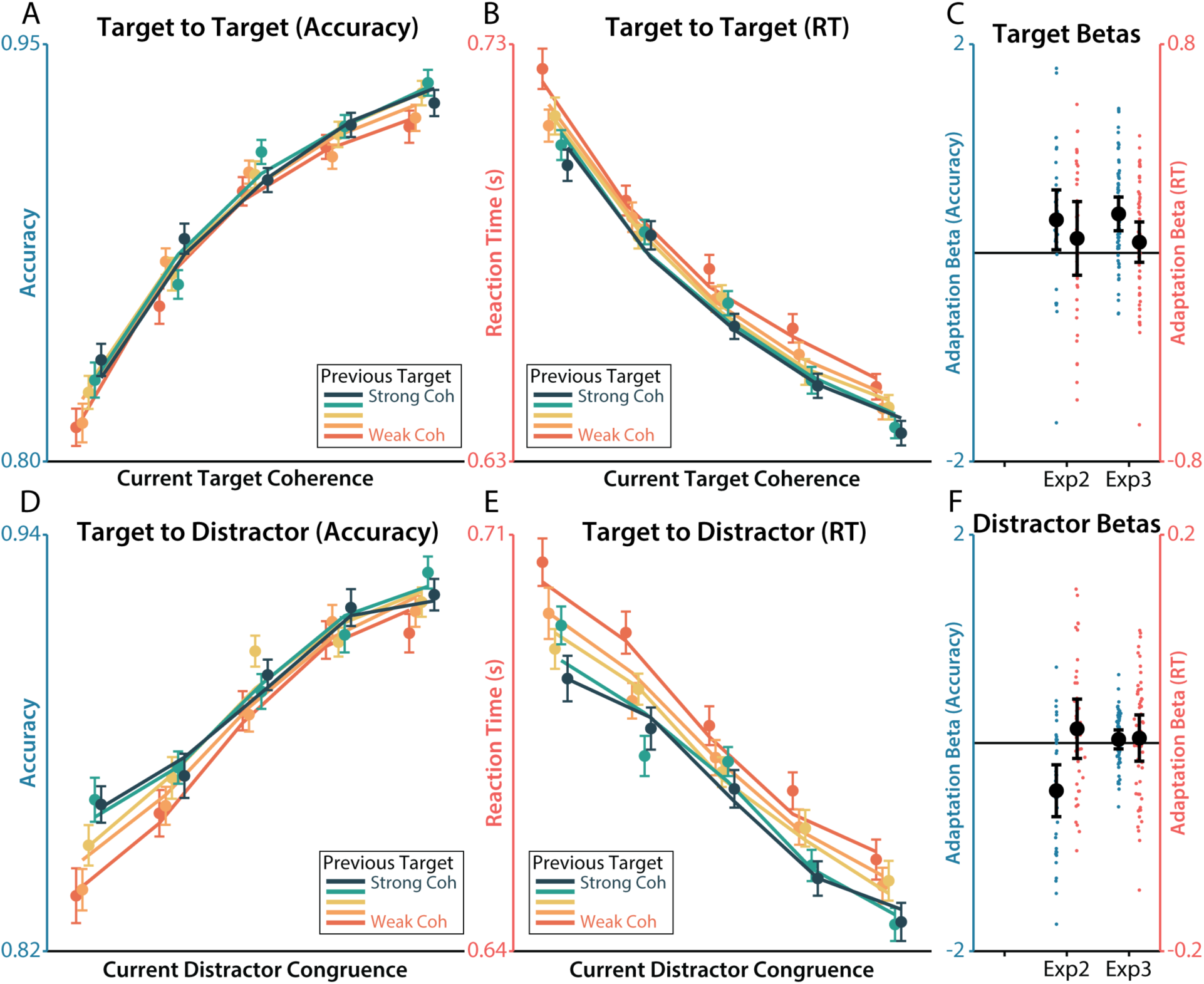
*Target-dependent adaptation.* **A-B)** The relationship between target coherence and accuracy (A) was weaker when the previous trial had weaker target coherence (redder colors). There was not a significant effect for RT (B) Lines depict aggregated regression predictions. **C)** Regression estimates for the current target coherence by previous target coherence interaction, within each experiment. **D-E)** There was not a significant relationship between distractor congruence and previous target coherence in accuracy (D) or RT (E). **F)** Regression estimates for the current distractor congruence by previous target coherence interaction, within each experiment. Error bars on behavior reflect within-participant SEM, error bars on regression coefficients reflect 95% CI.

**Supplementary Figure 3. (supplement to Figure 9).**
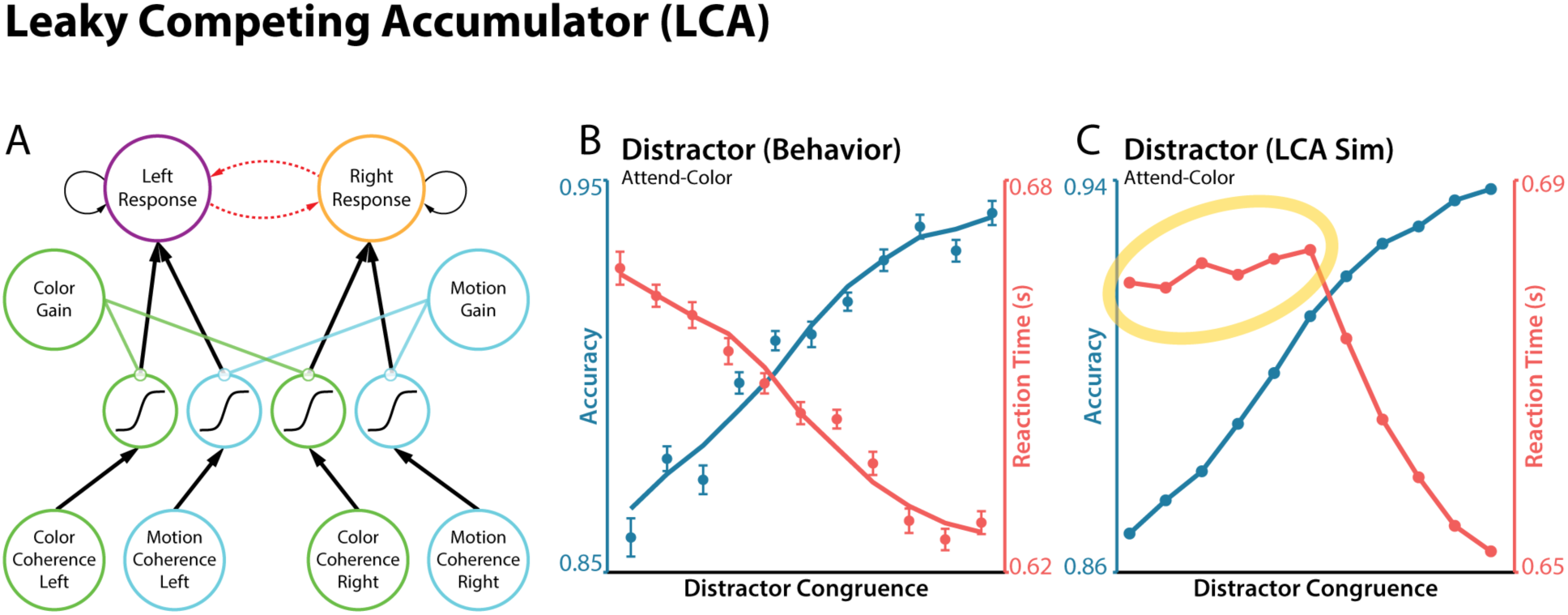
*Leaky competing accumulator simulation.* **A)** We simulated behavior from a leaky competing accumulator (Usher and McClelland, 2001). In this model, the response accumulators directly compete. In our parameter regime, leak and competition parameters produce race-like accumulation dynamics (Bogacz et al., 2006; Weichart et al., 2020). **B-C)** We found that this parameter regime was unable to capture the effect of distractor congruence on reaction time, as stronger inputs (congruent or incongruent) produce faster RTs in a race-like regime (Teodorescu and Usher, 2013). Other parameter regime, producing DDM-like dynamics, would replicate our main simulation results (Bogacz et al., 2006).

**Supplementary Figure 4. (supplement to Figure 10).**
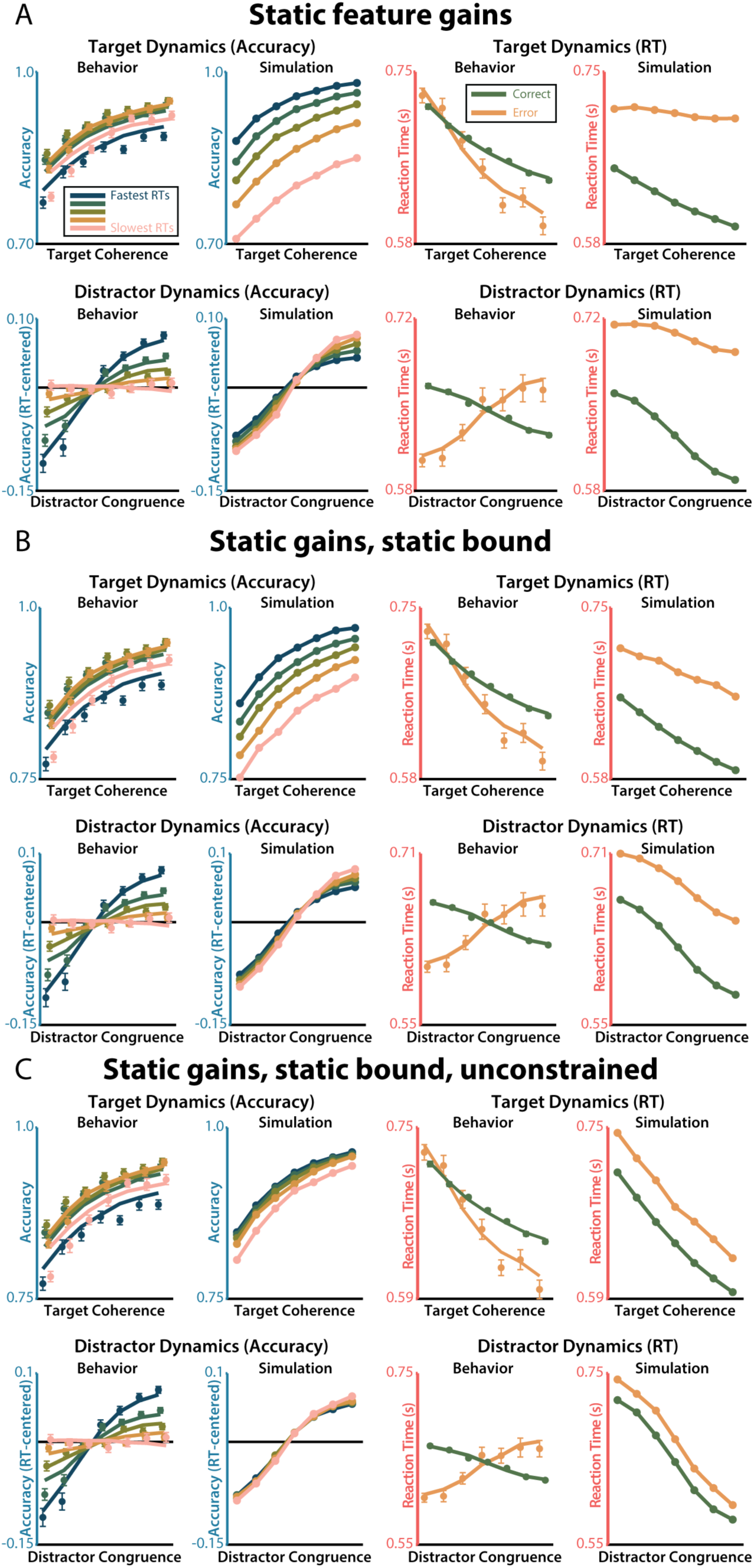
*Static feature gain simulations*. We simulated the FFI model under different formulations that lack feature sensitivity dynamics, showing that gain dynamics are necessary to capture the RT- and Accuracy-dependent feature sensitivity we observed in participants’ behavior. Feature-specific processes are necessary to capture the opposite-going dynamics on target sensitivity and distractor sensitivity. A) Static model without feature dynamics. B) Static model without feature dynamics or collapse response threshold. C) Static model without feature dynamics, collapsing response threshold, or positive-rectified accumulators.

**Supplementary Figure 5. (supplement to Figure 10).**
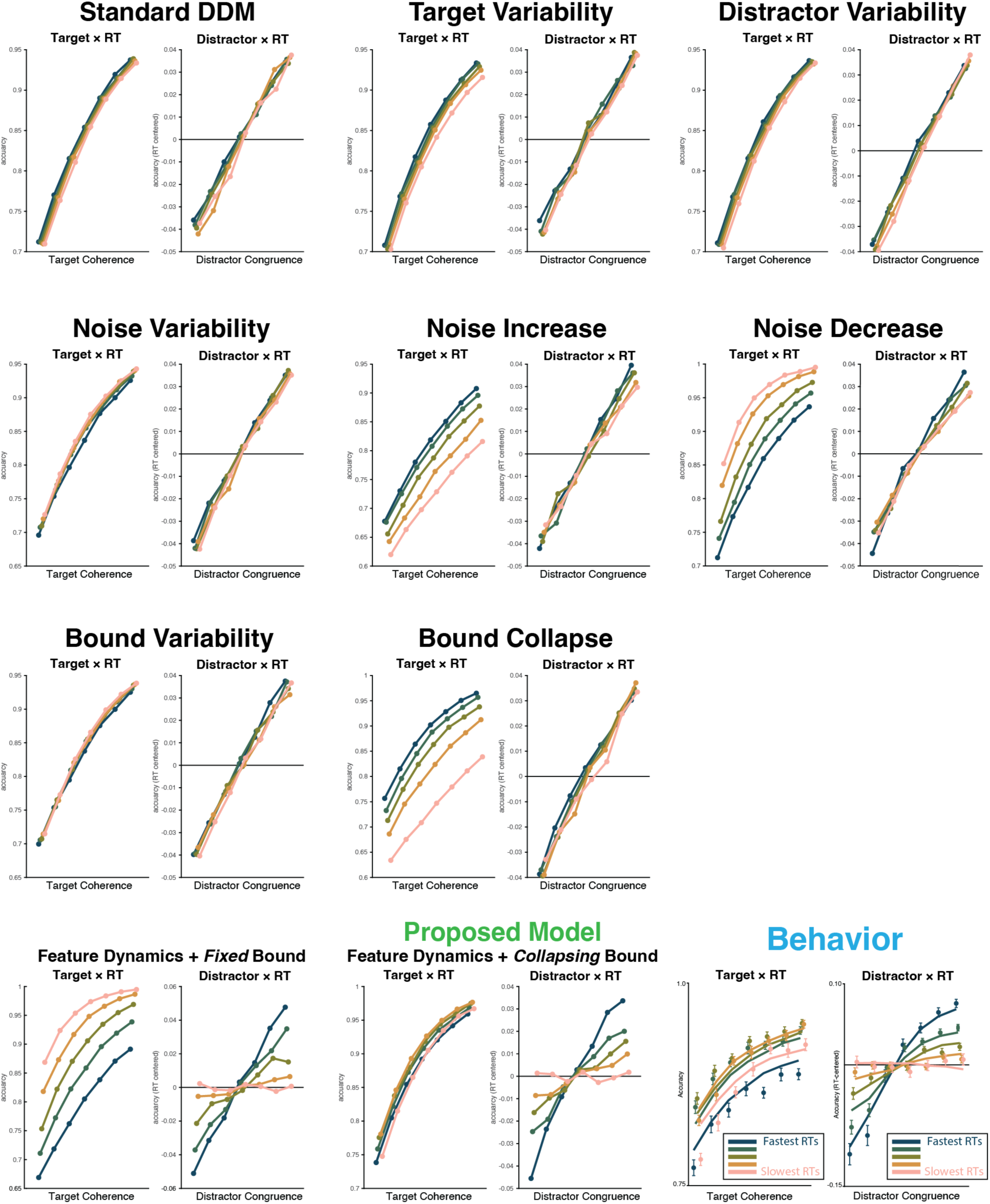
*Dynamic drift diffusion simulations*. Drift diffusion model (DDM) simulations demonstrating the predictions from alternative formulations of within- and across-trial dynamics. Data are simulated target and distractor psychometric curves, conditioned on simulated RT quintiles (1 million simulations per analysis). Row 1: Standard DDM, across-trial target gain variability, across-trial distractor gain variability. Row 2: across-trial accumulation noise variability, within-trial noise increase, within-trial noise decrease. Row 3: across-trial bound (threshold) variability, within-trial bound decrease (‘collapsing bound’). Row 4: within-trial target gain enhancement and distractor suppression with fixed bound, within-trial target gain enhancement and distractor suppression with collapsing bound, participants’ behavior. All simulations were performed using the dm package (package available at www.github.com/DrugowitschLab/dm; simulation scripts available at www.github.com/shenhavlab/PACT-public).

**Supplementary Figure 6. (supplement to Figure 2).**
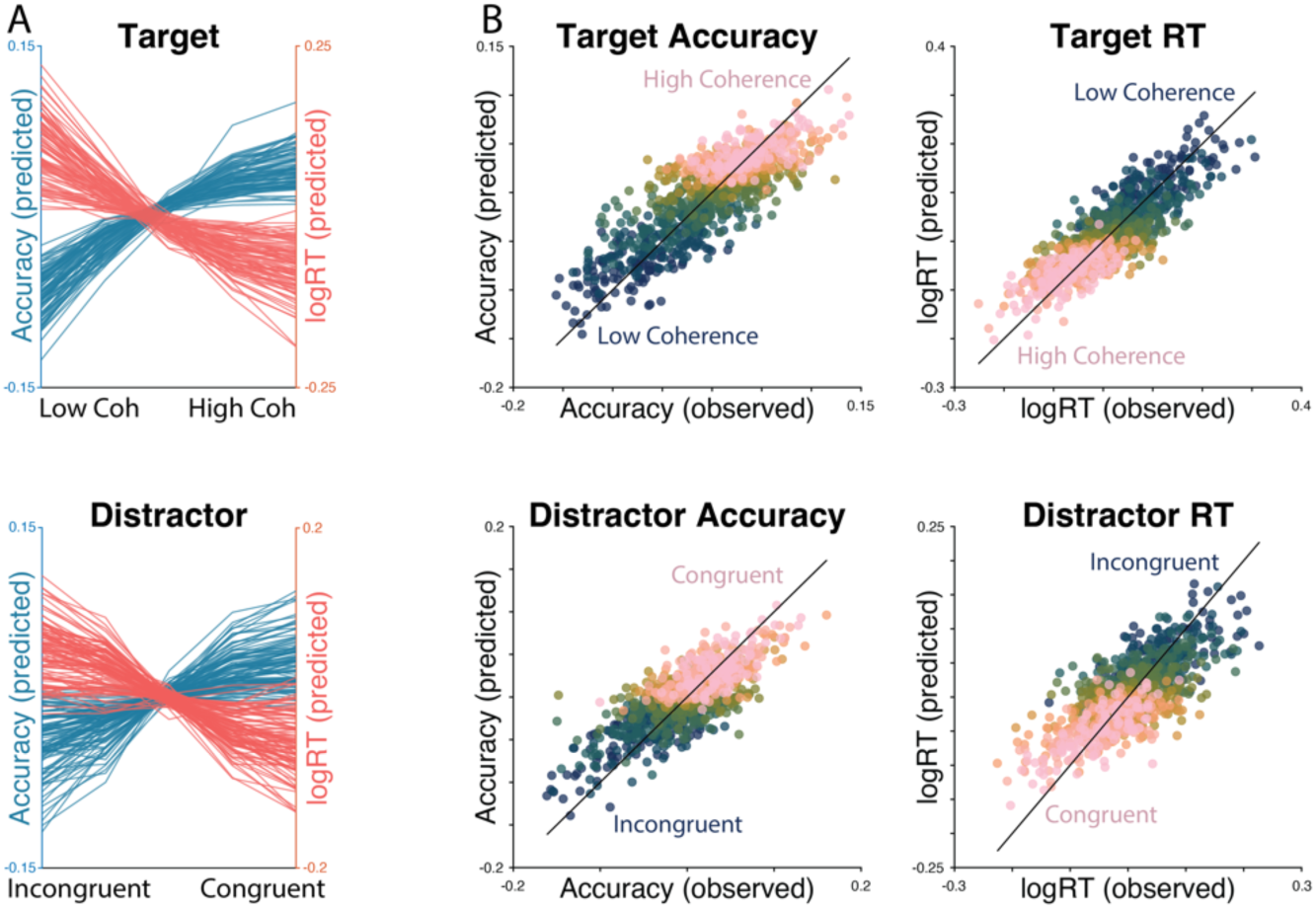
*Aggregated posterior predictive checks.* **A)** Model predictions from participants in Experiments 2 and 3, showing predicted target sensitivity curves (top) and distractor sensitivity curves (bottom). Predictions are centered within-participant to remove individual intercepts. **B)** Model fit quality for participants in Experiments 2 and 3. Each participants’ behavior (x-axis) is plotted against predicted behavior (y-axis), across five levels of target coherence (top) or distractor congruence (bottom; bluer to pinker indicates harder to easier conditions). Dots closer to the black identity reflect better model fit, and color gradients on y-axis reflect feature sensitivity. Predictions and behavior are centered within-participant to remove individual intercepts.

**Supplementary Figure 7. (supplement to Figure 2).**
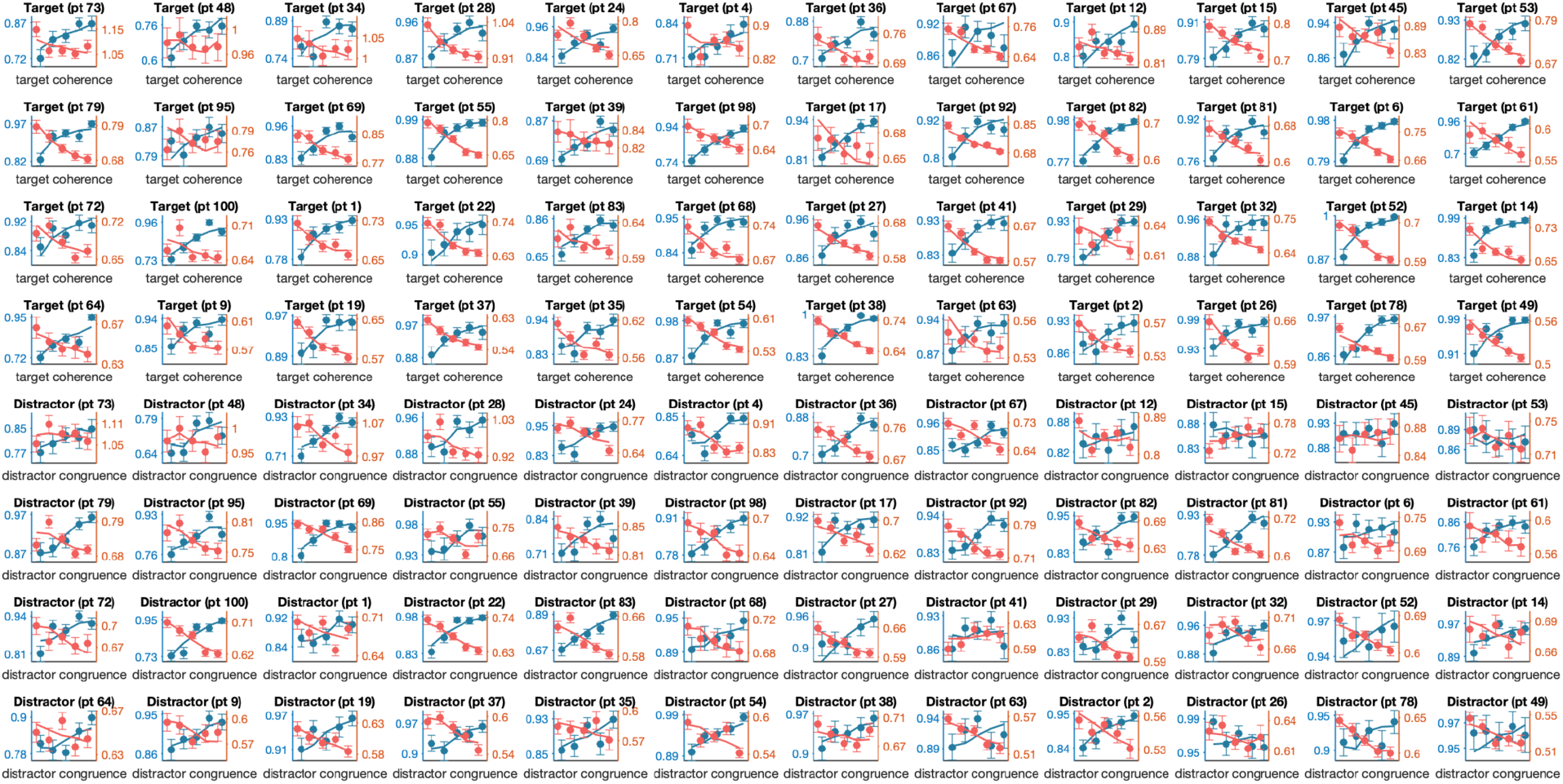
*Single-participant posterior predictive checks.* Posterior predictive checks from 48 participants from Experiments 2 and 3, linearly spaced from the poorest model likelihood to the best model likelihood. First four rows are target sensitivity curves for accuracy (blue) and reaction time (red). Final four rows are distractor sensitivity curves (for the same participants) for accuracy (blue) and reaction time (red). Overlaid lines are single-trial model predictions aggregated like participants’ behavior.

## Acknowledgements

This work was supported by the Daniel Cooper Graduate Student Fellowship (H.R.), as well as grants R01MH124849 and NSF CAREER Award 2046111 (A.S.). We are grateful to Kia Sadahiro, William McNelis, Allison Loynd, and Savannah Doelfel for assistance in data collection, and to Michael J. Frank, Matthew N. Nassar, Jonathan Cohen, David Badre, Tobias Egner, Senne Braem, Sebastian Musslick, and both the Shenhav Lab and LNCC for helpful discussions on these topics.

## Data Availability

Data and analysis scripts are available at github.com/shenhavlab/PACT- public

